# Vitamin auxotrophies shape microbial community assembly in the ocean

**DOI:** 10.1101/2023.10.16.562604

**Authors:** Rachel Gregor, Gabriel T. Vercelli, Rachel E. Szabo, Matti Gralka, Ryan C. Reynolds, Evan B. Qu, Naomi M. Levine, Otto X. Cordero

## Abstract

Microbial community assembly is governed by trophic interactions that mediate the transfer of carbon sources and biomass building blocks between species. However, central metabolism corresponds to only a small fraction of the biosynthetic potential of microbes: metabolites such as antimicrobial compounds, signaling molecules, and co-factors are underexplored forces shaping microbial communities. Here, we focus on B vitamin exchange in coastal marine bacterial communities that degrade particulate organic matter and find that natural seawater communities are vitamin limited. While almost a third of bacterial isolates from these communities are B vitamin auxotrophs, the pioneering degraders that first arrive on particles are vitamin producers that likely support auxotrophs through cross-feeding. However, combining experiments and a resource-explicit model, we show that auxotroph growth is often not restored by coculture with vitamin producers, but rather requires lysis and subsequent vitamin recycling. Our results highlight the importance of vitamin auxotrophies and lysis-mediated cross-feeding as important factors controlling microbial community assembly and succession on marine particles.

## Introduction

In the vast ocean, particles of organic matter serve as hubs of microbial activity and interactions.^1,2^ These particles, termed marine snow, are colonized and degraded by specialized bacteria as they sink, releasing sugars and other substrates like organic acids that support the growth of cross-feeding bacteria.^3,4^ Particle-associated communities harbor a dense network of interactions, through quorum sensing,^5^ antibiotics,^6^ and horizontal gene transfer.^7^ Metabolic dependencies frequently evolve in these communities, for example through the loss of genes to produce siderophores^8^ or hydrolytic enzymes.^9^ More broadly, metabolic dependencies such as vitamin auxotrophies are predicted to be prevalent among marine microbes^10,11^ and have been characterized in some marine isolates.^12–14^

It remains unclear how chemical interactions beyond the cross-feeding of sugars and organic acids shape marine bacterial community assembly. Vitamins are uniquely suited as a case study for chemical interactions, as these metabolites are highly conserved and required only in trace amounts, as coenzymes for central metabolic reactions. Vitamin exchange is important in algal-bacterial interactions^15^ and has been explored in other biomes,^16–18^ but has not been studied in the marine particle context. While some seawater amendment studies have shown that vitamins are limiting,^13,19^ it has also been proposed that trace vitamin levels in seawater are sufficient for auxotroph growth.^13,20^ If vitamins are indeed limiting, then biotic vitamin exchange is likely to be an important driver in community dynamics. This is especially relevant in the resource-rich particle environment, where other nutrients may not be limiting.

Here, we examine the role of vitamin exchange in the assembly of marine particle-associated bacterial communities. We combine natural seawater experiments with a large-scale characterization of vitamin requirements in 150 particle-associated isolates. We conclude that vitamin auxotrophies are important metabolic dependencies shaping particle community structure, and that vitamin cross-feeding may primarily take place through cell lysis.

## Results

### Vitamins are a limiting nutrient in coastal seawater microcosms

To determine if vitamins are limiting in particle-associated communities, we examined the effect of vitamin supplementation on coastal seawater microcosms incubated with model particles. As a simplified model for natural organic matter particles, we used particulate chitin, an abundant marine polysaccharide. We have previously shown that chitin particles are colonized by marine bacteria from the surrounding seawater in successive waves of degraders and cross-feeders.^3,4^ To test whether vitamins play a role in structuring these complex microbial communities, we incubated coastal seawater from Nahant, MA with chitin particles and supplemented half of the samples with a mixture of vitamins formulated for marine culture medium (Table 1; see Methods for details). Every 12 hours, we evaluated the impact of vitamins on community assembly by harvesting the particle-associated bacterial community and sequencing the samples using shotgun metagenomics.

**Table 1.** Vitamin concentrations in supplemented growth conditions.

| **Vitamin** | **Abbreviation** | **Final conc. (M)** |
| --- | --- | --- |
| Lipoic acid | ALA | $4.85 \times 10^{-7}$ |
| Thiamine hydrochloride | B1 | $2.97 \times 10^{-7}$ |
| Thiamine pyrophosphate | B1-PP | $2.17 \times 10^{-7}$ |
| Riboflavin | B2 | $2.66 \times 10^{-7}$ |
| Nicotinic acid | B3 | $8.12 \times 10^{-7}$ |
| Nicotinamide adenine dinucleotide (NAD) | B3-NAD | $1.51 \times 10^{-7}$ |
| Calcium d-pantothenate | B5 | $4.20 \times 10^{-7}$ |
| Pyridoxine hydrochloride | B6 | $4.86 \times 10^{-7}$ |
| D-Biotin | B7 | $1.23 \times 10^{-7}$ |
| Folic acid | B9 | $2.27 \times 10^{-7}$ |
| 4-aminobenzoic acid (PABA) | B9-PABA | $7.29 \times 10^{-7}$ |
| L-ascorbic acid | C | $5.68 \times 10^{-7}$ |
| Cyanocobalamin | B12 | $7.38 \times 10^{-9}$ |

We found that adding vitamins drastically altered the community composition and functional potential of seawater microcosms, indicating that these communities are indeed vitamin limited (Fig. 1). Key taxonomic groups that typically dominate chitin communities, such as *Vibrionales*, *Alteromonadales*, and some *Pseudomonadales,* reached a higher relative abundance in the absence of vitamins (Fig. 1a, Extended Fig. 1). This indicates that these taxa may have a competitive advantage in this condition and are likely vitamin producers (termed prototrophs). In vitamin-supplemented samples, taxa including *Rhodobacterales* and *Flavobacteriales* were enriched, suggesting that these may be vitamin auxotrophs (Fig. 1a). These taxonomic shifts led to differences in functional potential: vitamin-supplemented communities had a lower fraction of chitin degraders, as inferred from the frequency of chitin-hydrolysis protein domains in the metagenomes (Fig. 1b).

**Fig. 1.**
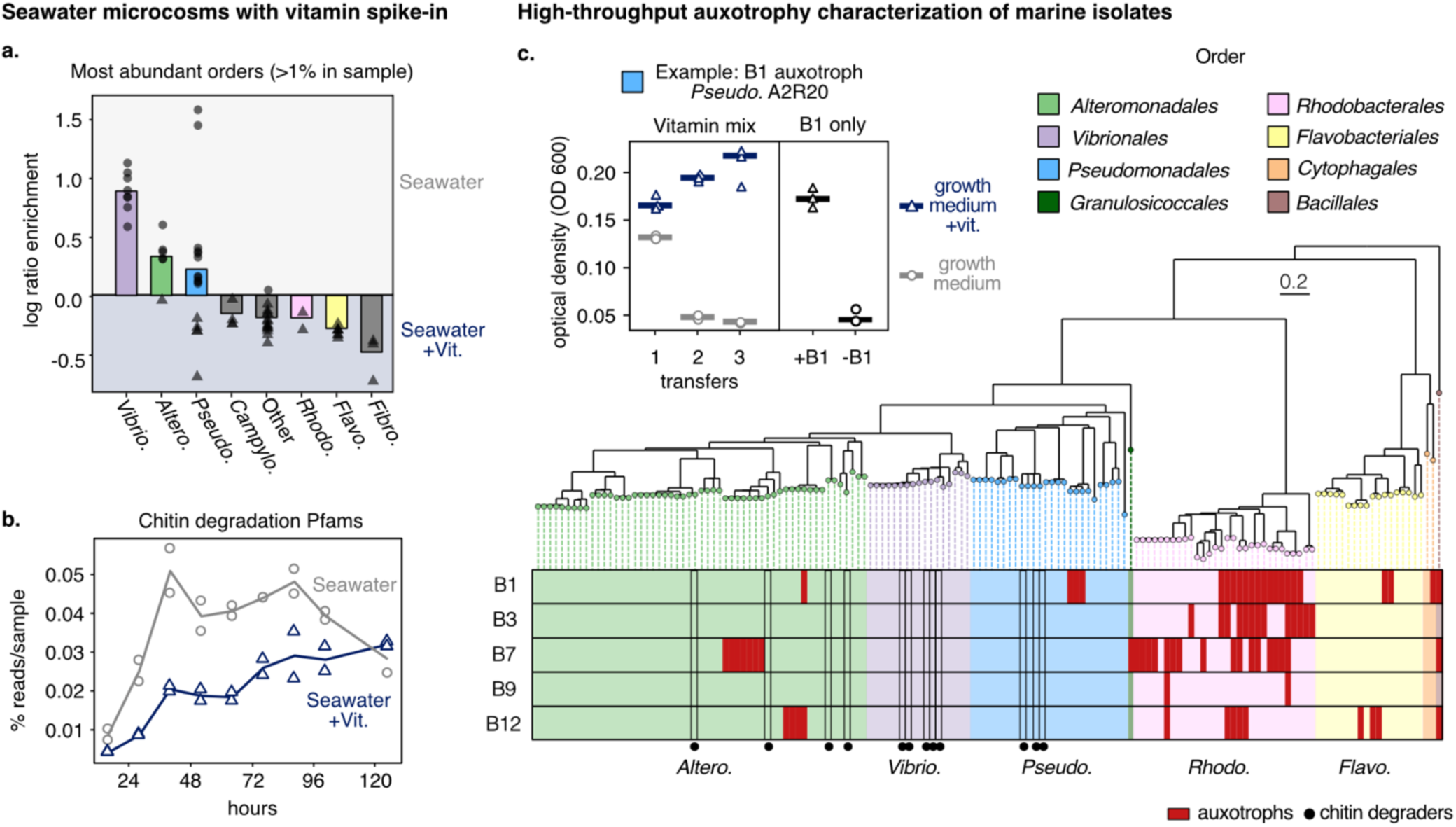
Coastal seawater communities are vitamin-limited and vitamin auxotrophies are prevalent. A. Coastal seawater collected at Nahant, MA, was incubated with chitin particles, with and without a vitamin mix spike-in. The addition of vitamins led to differences in taxonomy at the order level. Each data point represents a taxon that reached at least 1% abundance in one or more time points. B. Chitin degradation proteins families (Pfams) were depleted in the vitamin-supplemented seawater condition compared to the seawater condition. C. 150 bacterial coastal marine isolates from Nahant, MA, were screened for vitamin auxotrophies in a high-throughput format. Top left, example of growth with and without vitamin mix over three daily transfers, due to auxotrophy for B1. Chitin degraders were prototrophic (black circles). Abbreviations: *Vibrio*. = *Vibrionales*; *Altero*. = *Alteromonadales*; *Pseudo*.= *Pseudomonadales*; *Rhodo*. = *Rhodobacterales*; *Flavo*. = *Flavobacteriales*. *B-vitamin auxotrophies are prevalent in marine particle-associated isolates*

### B-vitamin auxotrophies are prevalent in marine particle-associated isolates

To understand the mechanisms driving the community response to vitamins, we characterized vitamin auxotrophies in a collection of coastal, particle-associated strains previously isolated from the same location (Nahant, MA).^21^ We characterized 150 seawater isolates representative of five of the orders affected by vitamin supplementation (Fig. 1c). To identify auxotrophies, we grew the isolates in a defined growth medium with and without vitamin supplementation (Table 1). We measured growth over three daily growth-dilution cycles, until vitamins from the initial seed cultures were diluted to picomolar concentrations, similar to environmental levels. Isolates with reduced growth without vitamins (Extended Fig. 2) were then assessed for growth with and without each vitamin individually to identify specific auxotrophies (Fig. 1c, top left).

**Fig 2.**
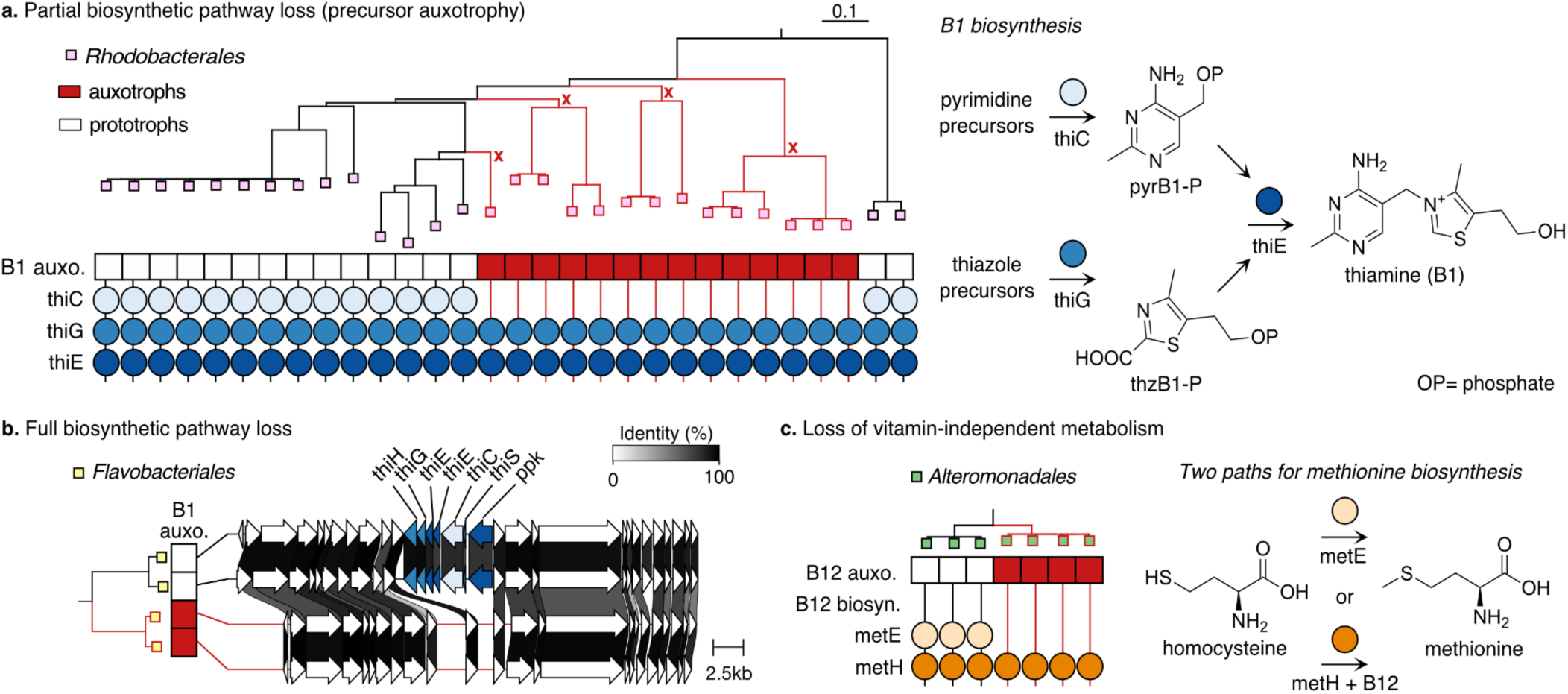
Three evolutionary modes of auxotroph formation. A. Partial pathway loss leading to formation of B1 pyrimidine auxotrophs in *Rhodobacterales*. Auxotrophies evolve frequently, with four putative loss events shown here (red “x”). Right, three critical genes in the thiamine biosynthesis pathway: *thiC*, which forms the pyrimidine precursor 4-amino-5-hydroxymethyl-2-methylpyrimidine phosphate (pyrB1-P); *thiG*, which forms the thiazole precursor 2-(2-carboxy-4-methylthiazol-5-yl)ethyl phosphate (thzB1-P); and *thiE*, which couples the two moieties to form thiamine. B. Total pathway loss. Genomic region of the thiamine biosynthesis cluster is lost in *Flavobacteriales* B1 auxotrophs compared to two closely related prototrophic isolates, with flanking regions conserved. C. Loss of vitamin-independent metabolism. Both prototrophic and auxotrophic *Alteromonadales* isolates are missing B12 biosynthesis genes and can scavenge B12. B12 auxotrophs have lost *metE*, retaining only the B12-dependent pathway for the biosynthesis of methionine via *metH*.

Consistent with the seawater microcosm experiment (Fig. 1a), all *Vibrionales* isolates were prototrophs, while 87% of the *Rhodobacterales* isolates were auxotrophic for one or more vitamins (Fig. 1c). These results support the hypothesis that the community response to vitamins in the seawater microcosms was likely driven by auxotrophies. Overall, almost one-third of the particle-associated isolates (47/150) were auxotrophic for vitamins in the supplementation mix (Fig. 1c). Further characterization revealed that the auxotrophies were for one or more of five key B vitamins: thiamine (B1), niacin (B3), biotin (B7), folic acid (B9), and cobalamin (B12).

We hypothesized based on the seawater microcosm data that degraders would be likely to be prototrophic, as chitin-hydrolysis genes were more abundant in the absence of added vitamins. Indeed, in a subset of 65 isolates characterized for growth on chitin,^4,9^ the 12 degraders were exclusively prototrophic, with representatives from the clades *Vibrionales*, *Alteromonadales*, and *Pseudomonadales* (Fig 1c, degraders marked with *). This was further supported by the isolate genomes, in which both chitin and alginate hydrolysis genes were enriched in prototrophs (Extended Fig. 3; Wilcoxon rank sum test: p-value < 0.001). This data suggests that vitamin prototrophy may provide a competitive advantage for degraders in particle-associated communities in the ocean.

**Fig. 3.**
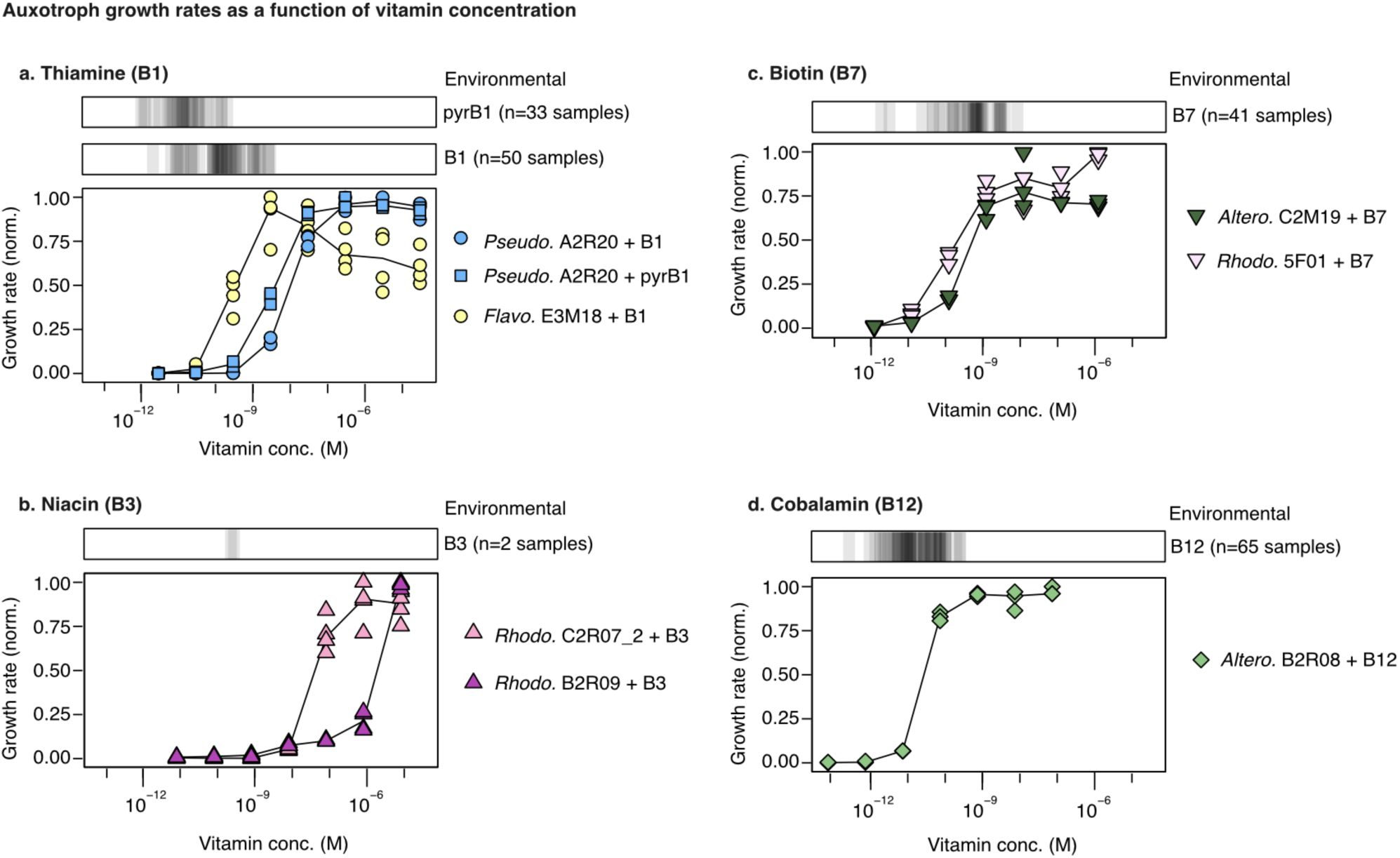
Auxotroph growth rates are limited under environmental vitamin concentrations. Auxotrophs were grown on different vitamin or precursor concentrations: A. thiamine (B1) and the precursor 4-amino-5-hydroxymethyl-2-methylpyrimidine (pyrB1), B. niacin (B3), C. biotin (B7), and D. cobalamin (B12). Cultures were grown for three growth-dilution cycles, diluting 1:40 every 24 hours. After the last transfer, a kinetic growth measurement was run and growth rates calculated from the exponential phase. Literature values for vitamin concentrations measured in the surface oceans are shown as gray bars above each panel (n=193 samples, Table S3).^26–53^ Abbreviations: *Altero*. = *Alteromonadales*; *Pseudo*.= *Pseudomonadales*; *Rhodo*. = *Rhodobacterales*; *Flavo*. = *Flavobacteriales*.

### Auxotrophies evolve through partial loss of biosynthesis pathways

We next established the genetic basis of vitamin auxotrophies by comparing the results of the phenotype characterization to the isolate genomes. We found a strong match between the observed auxotrophies and the absence of key vitamin biosynthesis genes based on previous studies^13,14,22–25^ (Extended Fig. 4, Table S1 and S2). Based on the presence-absence of these genes, the genomes were assigned as predicted prototrophs or auxotrophs. These predictions were generally accurate for B1, B3, and B7, with error rates between 3-7% (Table S2). This indicates that the observed auxotrophies were primarily the result of gene loss, as opposed to differential gene regulation or experimental error. However, in almost all cases only part of the biosynthetic pathway was lost, resulting in auxotrophies for vitamin precursors to feed into the remaining biosynthetic steps. For example, B1 auxotrophy in *Rhodobacterales* was driven by the loss of a key biosynthetic gene, *thiC*, resulting in an inability to synthesize the pyrimidine subunit of B1, although the rest of the pathway was retained (Fig. 2A). We observed fewer cases in which the entire biosynthetic pathway was lost, for example for B1 in *Flavobacteriales* (Fig. 2B). Notably, rare auxotrophies were less accurately predicted in this dataset: neither of the only two B9 auxotrophs in the collection were predicted correctly from their genomes.

**Fig. 4.**
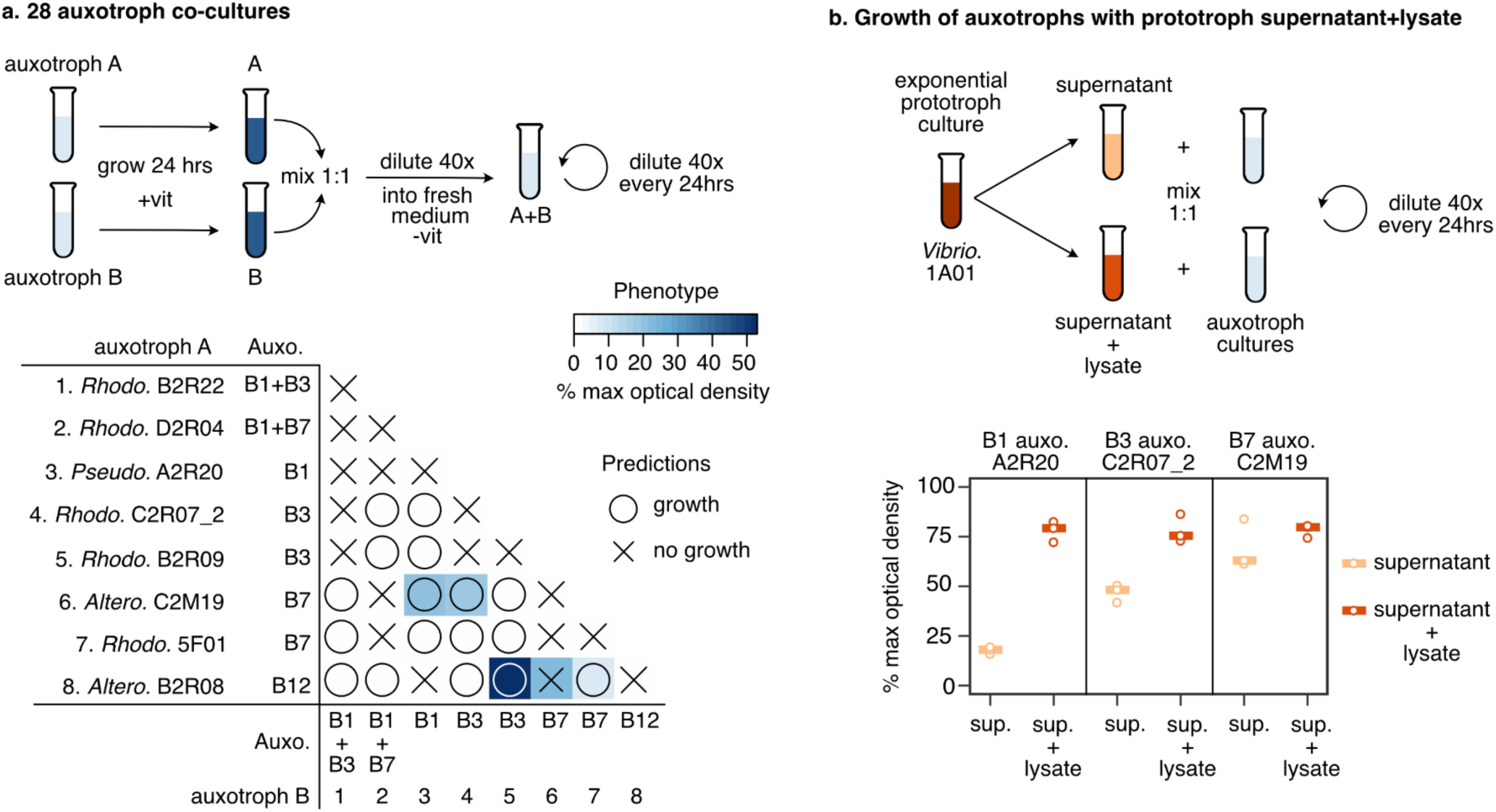
Auxotrophies are only partially alleviated in co-culture and by supernatants. A. Eight auxotrophs were grown in co-culture, for a total of 28 pairs. Top, the auxotrophs were grown separately with vitamins, then combined 1:1 and transferred to fresh medium without vitamins. Growth was measured after 3 growth-dilution cycles. Bottom, only 4/17 co-cultures predicted to successfully cross-feed (circles) grew to detectable levels, as well as 1/11 of the co-cultures predicted not to grow. The co-cultures reached between 7-52% of the final optical density of co-cultures with vitamin supplementation (% max optical density, blue heatmap). The 8 monocultures appear on the diagonal. B. Three auxotrophs (auxotrophs 3, 4, and 6) were further tested for growth with supernatants (peach) and lysates containing both supernatant and lysed cells (red), collected from a *Vibrio.* 1A01 prototroph culture in late exponential phase. Growth was measured after 3 growth-dilution cycles. All data is presented as a percentage of the optical density obtained when the supernatant is supplemented by vitamins (% max optical density), to correct for the effect of the depletion or release of additional nutrients in each condition (see Extended Fig. 8 for raw data). Abbreviations: *Vibrio*. = *Vibrionales*; *Altero*. = *Alteromonadales*; *Pseudo*.= *Pseudomonadales*; *Rhodo*. = *Rhodobacterales*.

By contrast, there was little correlation between vitamin B12 auxotrophies and predictions based on biosynthesis genes (Extended Fig. 4, Table S1 and S2). Vitamin B12 is not universally essential, as many bacteria and algae have alternate metabolic pathways that circumvent the need for a coenzyme. For instance, there are two canonical methionine biosynthesis pathways, of which only one requires B12 as a cofactor. Here, most of the isolates are predicted to scavenge B12 but do not produce or require it, and are therefore prototrophs. However, we observed one clade of *Alteromonadales* B12 auxotrophs that lost the alternate methionine biosynthesis pathway, leaving them with only the B12-dependent pathway (Fig. 2C). Only around 30 isolates in the collection were predicted to produce B12, primarily *Rhodobacterales* isolates with only the B12-dependent methionine biosynthesis pathway (Extended Fig. 5). Since the biosynthesis of B12 involves over 30 genes, B12 producers are commonly classified based on percent pathway completeness rather than specific genes.^25^ Here, this approach led to misclassifying some likely precursor auxotrophs with single gene losses as producers (Extended Fig. 5).

**Figure 5.**
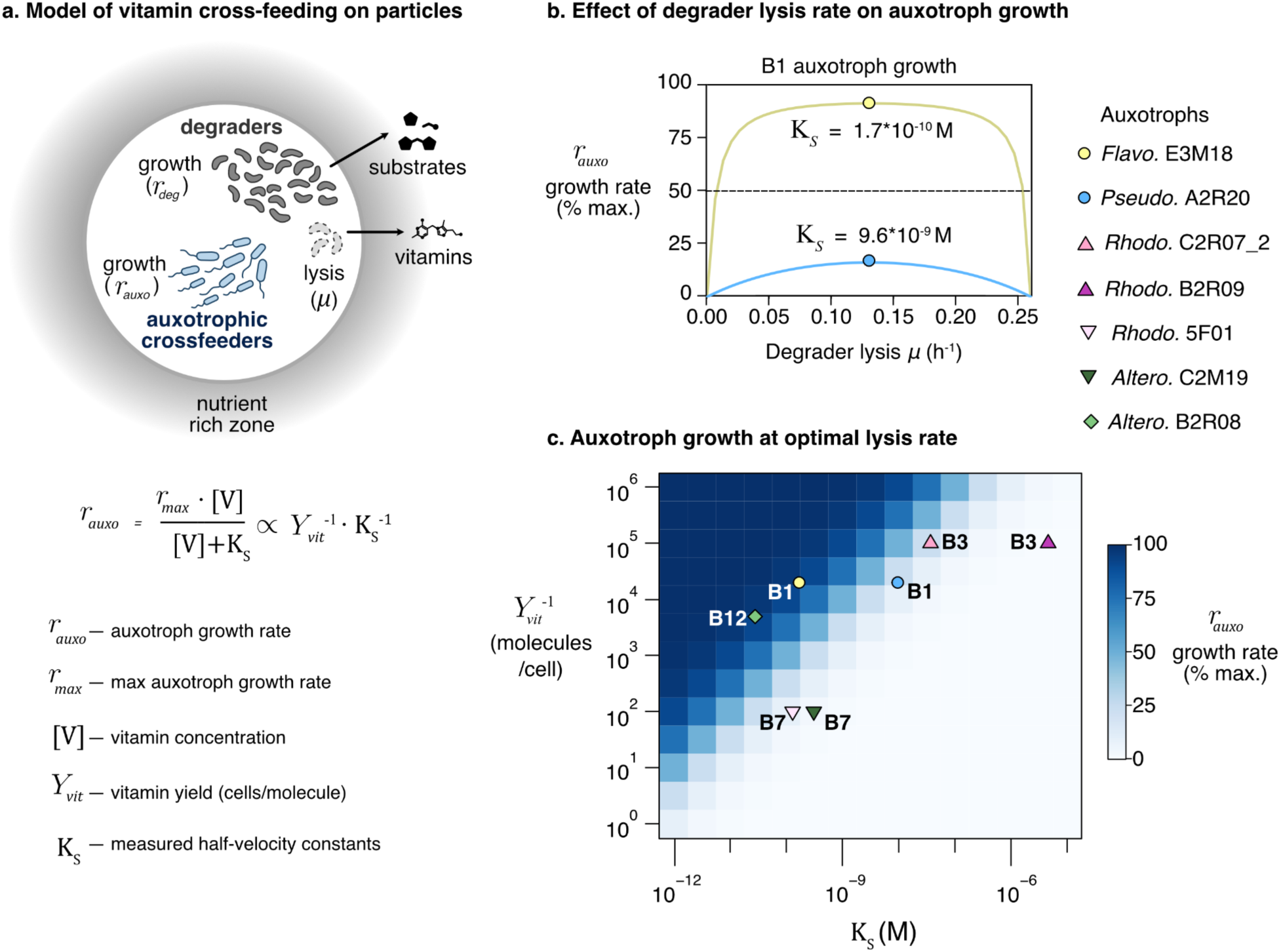
Model of vitamin cross-feeding in a particle-associated community. A. In a model particle community, the release of substrates is a function of the population size of degraders, which is set by their effective growth rate (*r_deg_*). The release of vitamins is assumed to be a function of the rate of degrader lysis (*μ*). The auxotroph growth rate (*r_auxo_*) depends on the vitamin concentration ([V]), half velocity constants (K_S_), and maximum growth rate for a given substrate concentration (*r_max_*), according to the Monod equation (bottom). Under diffusion and vitamin limitation, the auxotroph growth rate is approximately proportional to the inverse product of vitamin yield and affinity. (The inverse product is a generalizable quantity to determine growth outcomes in all conditions; see supplement for details.) B. Auxotroph growth rates increase rapidly as the degrader lysis rate increases, reaching a maximum at half the degrader growth rate. Two B1 auxotrophs with different half velocity constants reach 90% and 15% of their maximum growth rates (yellow circle and blue circle, respectively). C. Heatmap showing maximum auxotroph growth rate at optimal degrader lysis rates (blue shading), as a function of inverse vitamin yields (y-axis) and affinities (x-axis, half velocity constants K_S_). Vitamin auxotrophs in this study (symbols) achieve maximum growth rates ranging from approximately 0.2% to 95%.

Using the genome-based auxotrophy predictors, we expanded our analysis and found that the positive correlation between vitamin prototrophy and polysaccharide degradation is more broadly generalizable. We annotated over 11,000 diverse reference genomes from proGenomes^21^ and assigned auxotrophies for vitamins B1, B3, B7, and B9. On average, vitamin prototrophs had a higher number of glycoside hydrolases (polysaccharide-degrading enzymes) than auxotrophs, even after normalizing by genome size (Extended Fig. 6). Genomes with all four vitamin auxotrophies tended to have the fewest glycoside hydrolases. This suggests that polysaccharide degraders and vitamin auxotrophs may occupy separate ecological niches across environments.

### Auxotroph growth rates are limited under environmental vitamin concentrations

How do vitamin auxotrophs survive in the open oceans? The concentration of vitamins varies widely in the coastal oceans and is undetectable (sub-picomolar levels) in many areas.^26^ Since vitamins are required in low amounts and can be recycled, it has been proposed that amounts sufficient for survival can be scavenged from seawater. Alternatively, vitamin auxotrophs may require cross-feeding from more direct interactions with vitamin producers. The likelihood of either strategy depends on the amount of vitamins required to sustain growth. To determine growth requirements for vitamins, the growth of auxotrophs was measured across vitamin concentrations spanning seven orders of magnitude. We compared these values to environmental measurements of dissolved vitamins (n=193 samples, Table S3).^26–53^

The vitamin requirements of auxotrophs ranged from approximately 30 pM for a B12 auxotroph, to 5 μM for a B3 auxotroph (Fig. 3 and Table 2). While the two B7 auxotrophs had similar requirements, there was variation between auxotrophs for both B1 and B3. We hypothesized that this is due to different affinities for vitamins and their precursors, as has been shown in SAR11.^12^ However, a *Pseudomonadales* auxotroph had similar affinity for B1 and a B1 precursor, both an order of magnitude lower than a *Flavobacteriales* isolate (Fig. 3, top left). It is possible that this difference stems from variation in transporters or physiology and may vary for additional precursors.

**Table 2.**
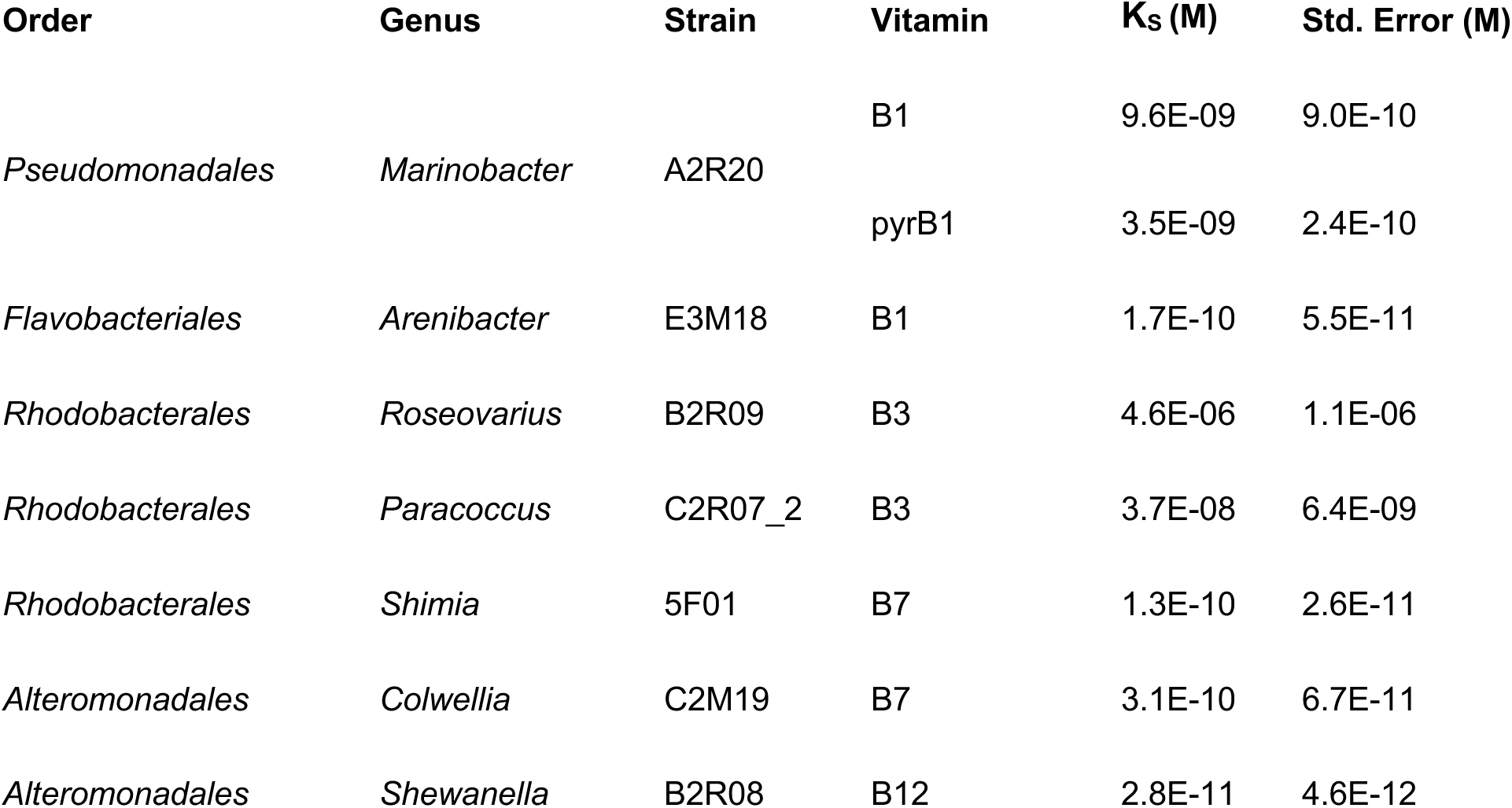
Estimated half-velocity constants (K_S_)

| Order | Genus | Strain | Vitamin | $K_s$ (M) | Std. Error (M) |
| --- | --- | --- | --- | --- | --- |
| *Pseudomonadales* | *Marinobacter* | A2R20 | B1 | 9.6E-09 | 9.0E-10 |
|  |  |  | pyrB1 | 3.5E-09 | 2.4E-10 |
| *Flavobacteriales* | *Arenibacter* | E3M18 | B1 | 1.7E-10 | 5.5E-11 |
| *Rhodobacterales* | *Roseovarius* | B2R09 | B3 | 4.6E-06 | 1.1E-06 |
| *Rhodobacterales* | *Paracoccus* | C2R07_2 | B3 | 3.7E-08 | 6.4E-09 |
| *Rhodobacterales* | *Shimia* | 5F01 | B7 | 1.3E-10 | 2.6E-11 |
| *Alteromonadales* | *Colwellia* | C2M19 | B7 | 3.1E-10 | 6.7E-11 |
| *Alteromonadales* | *Shewanella* | B2R08 | B12 | 2.8E-11 | 4.6E-12 |

The growth rates of all auxotrophs were limited under the majority of measured environmental vitamin concentrations in seawater (Fig. 3, gray bars; Table S3). This was especially stark for vitamin B3, which was recently quantified at approximately 50 pM in seawater,^51,52^ 3-5 orders of magnitude lower than the requirements measured here. While far fewer measurements exist for particulate fractions, vitamin levels are similar to or even lower than in dissolved seawater^44,52^ (Table S3). This data confirms that auxotroph growth is likely often vitamin-limited in the global oceans, especially in nutrient-rich particle communities where other resources are at high local concentrations. The striking similarity between the environmental concentrations and the measured bacterial affinities suggests that bacterial uptake may dictate vitamin levels in parts of the oceans, by depleting them to the lowest concentration possible.

### Auxotroph co-cultures are vitamin-limited

If ambient vitamin levels in seawater are not sufficient for growth, vitamin auxotrophs likely rely on cross-feeding with vitamin producers. Cross-feeding in co-culture has been shown with algae,^15,54,55^ with engineered vitamin auxotrophs,^56,57^ and for organic acids with isolates from this collection.^3,58^ We therefore hypothesized that natural bacterial auxotrophs would complement one another’s vitamin requirements during growth. To test this, we grew a selection of eight auxotrophs in co-culture, including two dual auxotrophs. Each auxotroph was initially grown in monoculture to high density in vitamin-supplemented media then combined in pairs (Fig. 4a, top). We measured co-culture growth after three growth-dilution cycles, diluting 1:40 every 24 hours, and compared it to growth in vitamin-supplemented media. Of the 28 co-cultures, 17 pairs were predicted to be able to successfully complement one another’s vitamin auxotrophies, while 11 pairs were predicted not to grow.

Most auxotroph pairs did not grow in co-culture (Fig. 4a). Only around one quarter (4/17) of the predicted successful pairs grew to detectable levels, and that growth was vitamin-limited, ranging from just 7-52% of growth with vitamin supplementation. The extinction of most of these co-cultures indicates that they divided fewer than approximately 5 times over 24 hours (log_2_ of the dilution factor 40x). This is despite being initially combined in relatively high density after growth in vitamin-rich media to avoid Allee effects, i.e. limitations due to positive density-dependent growth^59^. While the two dual auxotrophs did not grow in any co-culture, the other six isolates each grew in at least one instance. Almost all of the 11 co-cultures predicted not to grow indeed did not, except for one: the *Altero.* auxotrophs for B7 (C2M19) and B12 (B2R08). While C2M19 is unable to synthesize B12 based on its genome, it can produce methionine, which can alleviate the need for B12 in the *Altero.* B12 auxotrophs (Fig. 2c). Overall, this data indicates that even when cross-feeding is theoretically possible, most pairs of auxotrophs are not able to grow together in co-culture.

We next hypothesized that prototrophic degraders supply vitamins to auxotrophic cross-feeders (Fig. 4b). While a subset of auxotrophs grew with prototrophs in co-culture to varying ratios (Extended Fig. 7), prototroph growth rates were higher than auxotrophs and quickly reached stationary phase. We therefore could not differentiate between facilitation by secretion of vitamins during growth or by lysis in stationary phase. To further study the mechanism of facilitation, we selected the *Vibrio.* chitin degrader 1A01, which is predicted to produce vitamins B1, B3, and B7, and collected supernatants and lysates during late exponential growth (Fig. 4b, top). We tested three auxotrophs from the panel, growing them with the supernatants and lysates mixed 1:1 with fresh media (Fig. 4b, bottom). All data is presented as a percentage of growth in supernatants with added vitamins, to represent the maximal possible growth given the depletion and release of other nutrients (see Extended Fig. 8 for raw data).

Auxotroph growth was partially restored by prototroph supernatants and more so by lysates, depending on the vitamin (Fig. 4b, bottom). All three auxotrophs reached approximately 78% of maximal growth when grown on lysates, which contain both supernatants and lysed cells (Fig. 4b, red). The growth on supernatants was variable for different vitamins, ranging from only 18% of maximal growth for B1 auxotroph *Pseudo.* A2R20 to 69% for B7 auxotroph *Altero.* C2M19 (Fig. 4b, peach). The extent of facilitation also depends on the physiology of the prototroph, with different trends for each vitamin in early exponential and stationary phase cultures (Extended Fig. 8). This data indicates that the externalization of vitamins during growth is highly variable even for a single vitamin prototroph and raises the possibility that lysis may be an important mechanism in facilitating cross-feeding.

### Low lysis rates alleviate vitamin auxotrophies on particles

Metabolites are rapidly lost to diffusion in the open oceans. Under these conditions, cross-feeding requires high metabolite concentrations and/or high uptake affinities, even in densely packed and spatially structured particle communities. Given these constraints, we asked whether lysis can sustainably support vitamin cross-feeding on particles without leading to community collapse. To this end, we developed and parameterized a spatial model for the growth and lysis of degraders and auxotrophs on a particle, accounting explicitly for the production, diffusion, and uptake of substrates and vitamins (Fig. 5a, Table S4; model details in SI text).

The model predicts that auxotroph growth rates quickly increase as degraders lyse, and then decrease as high lysis rates lead to a reduction and eventual collapse of the population (Fig. 5b). Under our assumptions, the lysis rate optimal for auxotroph growth is equal to half the maximum degrader growth rate (0.26 h^-1^), representing a decrease from 9 to 4.5 doublings a day. The extent of facilitation is highly dependent on the amount of vitamins released per lysed cell (yield), and the auxotroph’s vitamin affinity (half velocity constant) (Fig. 5a, bottom). For example, the B1 auxotroph *Flavo.* E3M18 quickly reaches half its maximum growth rate at a very low degrader lysis rate (0.007 h^-1^), and at optimal lysis rates reaches 90% maximum growth (Fig. 5b, yellow circle). In comparison, the B1 auxotroph *Pseudo.* A2R20 has a lower vitamin affinity, and even at an optimal lysis rate reaches only 15% of its maximum growth rate (Fig. 5b, blue circle).

The model predicts that degrader lysis can sustainably support growth on particles under diffusion for the majority of auxotrophs in this work (Fig 5c). In the parameter space of possible combinations of vitamin yields and affinities, the auxotrophs fall on a transition region that approximately follows the 50% growth line. This suggests some level of evolutionary fine-tuning for affinities that match vitamin requirements: higher affinities evolve to efficiently consume rare resources, and vice versa. Auxotroph growth rates span a wide range of values (blue shading), suggesting that auxotroph growth is sensitive to variations in vitamin yields, affinities, and local lysis rates. While we find that auxotroph growth from lysis is generally possible, two auxotrophs are predicted to grow extremely slowly, at less than 5% of their maximal growth rates (Fig. 5c, Table S5). Other mechanisms may explain the growth of these auxotrophs in nature, for example the secretion of vitamins and their precursors, as has been observed for B7.^14^

## Discussion

We found that B vitamin auxotrophies are prevalent in marine particle-associated communities. In coastal seawater microcosms, the addition of vitamins altered community composition and reduced the community chitin degradation potential (Fig 1). These effects were likely due to vitamin auxotrophies, which we found in almost a third of particle-associated isolates in a high-throughput characterization. Vitamin auxotrophies for B1, B3, and B7 are generally correctly predicted from genomes, and most often evolve through partial pathway loss (Fig 2). The growth rates of all auxotrophs were limited under even maximum dissolved environmental vitamin concentrations (Fig 3), leading us to hypothesize that vitamin requirements are met through interactions. However, when auxotrophs were grown in pairs, most co-cultures did not grow (Fig. 4a). Some auxotroph growth was supported by supernatants and lysates collected from a prototroph degrader, depending on the vitamin and growth phase (Fig. 4b).

Vitamin auxotrophy in particle-associated communities was found in cross-feeders, not degraders. We found that all of the degrader isolates in our collection were prototrophs (Fig. 1c), and that hydrolytic enzyme genes were more abundant in prototrophs than auxotrophs (Extended Fig. 3). This was further supported by the seawater microcosm data, in which chitinase genes were more abundant in samples without vitamin supplementation (Fig. 1b). These degraders may also play an additional role as vitamin providers to auxotrophic cross-feeders, as is suggested by our supernatant experiments (Fig. 4b). This data fits well with previous findings that degraders are the first to arrive on particles, followed by a wave of cross-feeders.^60^ It may therefore be more advantageous for pioneering degraders to be self-sufficient prototrophs than for cross-feeders, which are reliant on other community members regardless. We observed the same pattern in a broad analysis of vitamin auxotrophies and polysaccharide degradation genes in over 11,000 genomes, indicating that this division into separate ecological niches may be more broadly applicable (Extended Fig. 6).

The patchy distribution of auxotrophies suggests that vitamin biosynthesis genes are frequently lost in particle-associated communities. In the majority of cases, only part of the biosynthetic pathway was lost, as has been recently observed in other vitamin auxotrophy studies in marine bacteria.^12–14^ This partial pathway loss leads to auxotrophies for vitamin precursors, and may also make it possible to regain vitamin biosynthesis through horizontal gene transfer. We conclude that vitamin biosynthesis is heterogeneously distributed in natural communities and that auxotrophies have evolved multiple times through independent loss events, as has been shown in algae^61^.

How are auxotrophs meeting their vitamin needs? We found that auxotroph growth was limited under environmental vitamin concentrations, and therefore likely requires cross-feeding with other bacteria. However, the majority of auxotrophs could not grow in co-culture, and growth was limited in most cases from prototroph supernatants as well (Fig. 4). Vitamin secretion during growth has been linked to overproduction and secretion via transporters.^62^ This data indicates that vitamin secretion is relatively rare and strain-specific, as has also been recently found in *Rhodobacterales* isolates for B12.^63^ Therefore vitamins are not necessarily shared public goods, in contrast to sugars released from particle degradation,^9^ excreted organic acids,^58^ or siderophores.^8^

If vitamins are only sometimes secreted during growth, a likely route for cross-feeding in particle communities is through cell lysis. Phage predation is estimated to kill 20-40% of bacteria in the surface oceans daily,^64–66^ and can facilitate cross-feeding,^67,68^ alter host metabolism,^69,70^ and cause metabolite release in particle communities.^71^ We modeled auxotrophic cross-feeder growth on particles under diffusion, and found that in most cases a relatively low rate of degrader lysis is sufficient to alleviate vitamin limitation (Fig. 5). Vitamin affinities are also negatively correlated with vitamin requirements, suggesting that like prices in a market, affinities are higher the rarer the commodity. Auxotroph growth on particles exists on a tight margin, sensitive to small changes in affinities, yields, and lysis or secretion rates.

The widespread prevalence of vitamin auxotrophies and likelihood of cross-feeding by lysis has two important implications. First, when studying bacteria in the lab, we almost always provide a surplus of nutrients such as vitamins. To translate these studies to the environment, we must carefully consider and study the effects of nutrient limitation on bacterial physiology and interactions. Second, cross-feeding by lysis is a general mechanism for metabolite exchange, agnostic to species identity. This is in direct contrast to specific species-species interaction networks based on pairwise co-cultures. The relative contribution of these two paradigms would greatly impact the flow of resources in natural microbial communities, and the extent to which microbial community composition is stochastic or predictable.

## Materials and methods

### Media and Growth Conditions

All experiments with isolates were performed at room temperature or 25 °C incubators in liquid culture using minimal media (MBL, see below). For co-cultures and kinetic experiments, frozen individual environmental isolates were streaked onto marine broth (Marine Broth 2216, Difco) agar plates, and single colonies were inoculated into marine broth liquid medium.

MBL media is a defined marine minimal media adapted from a protocol from the Marine Biological Laboratory Microbial Diversity course. The standard recipe contains the following components: 4x sea salt mix (1L water containing 80 g NaCl, 12 g MgCl_2_•6H_2_O, 0.60 g CaCl_2_•2H_2_O, 2.0 g KCl); 1000x trace mineral mix (1 L 20 mM HCl containing: 2100 mg FeSO_4_•7H_2_O, 30 mg H_3_BO_3_, 100 mg MnCl_2_•4H_2_O, 190 mg CoCl_2_•6H_2_O, 24 mg NiCl_2_•6H_2_O, 2 mg CuCl_2_•2H_2_O, 144 mg ZnSO_4_•7H_2_O, 36 mg Na_2_MoO_4_•2H_2_O; 25 mg NaVO_3_, 25 mg NaWO_4_•2H_2_O, 6 mg Na_2_SeO_3_•5H_2_O; note: NaVO_3_ and NaWO_4_•2H_2_O solids were handled in the chemical hood); 100x nitrogen source (1 M NH_4_Cl); 500x phosphorus source (0.5 M Na_2_HPO_4_); 1000x sulfur source (1 M Na_2_SO_4_); 20x HEPES buffer (1 M HEPES sodium salt, pH 8.2), and a carbon source as detailed below. All components were filter sterilized with 0.2 µm filters. Sea salt mix was stored at room temperature; sulfur, nitrogen, carbon, and phosphorus sources were stored in 4 °C; vitamin, trace metals, and HEPES were aliquoted into single use aliquots and frozen at −20 °C.

For the phenotype characterization, four carbon sources (glucose, glutamine, glycerol, and pyruvate) were added to the MBL media to support the growth of isolates with different nutrient preferences. Each carbon source was normalized by its number of carbon atoms to achieve a final concentration of 15 mM carbon each (2.5 mM glucose; 3 mM glutamine; 5 mM sodium pyruvate; 5 mM glycerol). For the kinetic growth curves as well as the supernatant/lysate experiments, a single carbon source (glucose for prototroph 1A01, pyruvate for auxotrophs) was used at 30 mM carbon.

### Seawater microcosms

Nearshore coastal seawater was collected from Nahant, MA on Oct 21, 2021, and filtered sequentially with 63 µm and 5 µm filters. Hydrogel chitin particles (New England Biolabs, #E8036L) were washed in sterile artificial seawater (Sigma-Aldrich, #S9883) and filtered through 100 µm mesh filters. The flowthrough was then passed through a 40 µm filter, and the particles captured on the filter were resuspended in artificial seawater. The particles were added to 50 mL aliquots of filtered seawater, for a final concentration of approximately 750 particles per tube (15 particles/mL). To half of the tubes, vitamins were added using the vitamin stock from the MBL media (Table 1). The tubes were rotated end over end at room temperature over five days. Every 12 hours, we harvested 3 replicate tubes from each condition and separated the chitin particles and associated bacterial community using a neodymium magnet. The seawater supernatant was removed with a serological pipette, leaving approximately 1 mL seawater and beads. The seawater+beads fraction was transferred to an Eppendorf tube and 0.5 mL of artificial sea salt solution was added. The samples were frozen at −20 °C.

Genomic DNA was extracted from 2 replicate tubes per condition using the Qiagen DNeasy Blood & Tissue Kit. The DNA yield was quantified using the Invitrogen™Quant-iT™ PicoGreen™ dsDNA Assay, with yields ranging between 0.01-0.04 ng/μl. The samples were processed and sequenced by the Microbial Genome Sequencing Center using the Illumina DNA Prep kit and protocol, modified for low concentrations. Paired end sequencing was run on an Illumina NextSeq2000, yielding 2×151bp reads. Raw sequencing reads were quality trimmed with Trimmomatic v0.36.^72^ The metagenomes were annotated for taxonomy using phyloFlash^73^ and using a custom database of profile hidden Markov models of proteins involved in growth on chitin^71^.

### Vitamin auxotrophy characterization

For a full list of strains, including their taxonomy and isolation details, see Supplementary File 1. Apart from the *Vibrionales* strains YB2^74^ and 1A06, 12B01 and 13B01^75^, all strains were isolated from enrichments of coastal seawater on polysaccharides^3,4,21^. A total of 187 strains were originally double-streaked the time of isolation and double-streaked again before being arrayed in two 96-well plates and stored in 5% dimethylsulfoxide at −80°C.

For the phenotype characterization, the arrayed isolate collection was defrosted and inoculated 1:10 into marine broth liquid medium in 1 mL 96-well plates. After 72 hours, the plates were transferred 1:100 into MBL medium. After 24 hours, the culture was transferred 1:40 into standard 96-well plates in two conditions, each in triplicate: standard MBL medium containing vitamins (MBL^+Vit^; see vitamin list in Table 1) and MBL without the addition of vitamins (MBL^-Vit^). Isolates that did not grow consistently upon being transferred to MBL^+Vit^ were dropped from the screen, leaving 150 isolates in total.

Every 24 hours, bacterial growth was estimated by measuring the optical density at 600 nm (OD_600_) using a TecanSpark plate reader with a stacker module. Then the plates were transferred 1:40 into fresh medium (MBL^+Vit^ or MBL^-Vit^). Plates were incubated at room temperature without shaking.

The isolates were classified as vitamin auxotrophs using the growth data from the third transfer (72 hours). The median OD_600_ value was calculated for both conditions and the growth deficit calculated as follows: (MBL^-Vit^ -MBL^+Vit^)/(MBL^-Vit^ + MBL^+Vit^). A growth deficit value of −0.4 was chosen as the cutoff for putative auxotrophs based on the distribution of values, with 52 isolates passing this threshold (Extended Fig. 2). The putative auxotrophs were then grown individually and assayed for growth with each vitamin separately, with supplementation of single vitamins as well as mixes of all vitamins but one. Each putative auxotroph was tested at least twice and the results compared. 47 of the putative auxotrophs were confirmed, with the remaining five removed as their phenotype was not consistent.

### Genome analysis

We established the genetic basis of vitamin auxotrophies by comparing the results of the phenotype characterization to the isolate genomes. The sequenced genomes^3,4,21^ were annotated using eggnog v4.5 (mmseqs mode with default parameters)^76^ and dbCAN v2 (diamond mode)^77^ for CAZymes. Taxonomy assignment and the creation of phylogenetic trees were performed using the standard workflow in GTDB-tk^78^, followed by renaming of taxa falling into the NCBI clades Vibrionales and Alteromonadales (both assigned Enterobacterales by GTDB-tk) to use the more familiar names for those clades^79^. The eggnog annotations were searched for vitamin biosynthesis genes based on previous studies^13,14,22–25^ (Extended Fig. 4, Table S1 and S2).

### Growth curves

To determine growth requirements for vitamins, the growth of auxotrophs were measured across vitamin concentrations spanning seven orders of magnitude. A selection of auxotrophs (Fig. 3) for different vitamins were chosen and grown in the same transfer protocol described above in the phenotype characterization, with two key changes: 1) Each auxotroph was initially grown in marine broth and then MBL^+vit^ as specified above, then transferred to eight conditions in total: MBL^-vit^, and seven ten-fold dilutions of a single vitamin in MBL^-vit^. The vitamin conditions were present in each transfer. 2) After the third transfer, the OD_600_ was measured continuously in the plate reader for 24-36 hours. The resulting growth curves were visualized and the growth rate calculated for each vitamin concentration from the exponential region using the lm function in R.

### Co-culture and supernatant experiments

We grew a selection of eight auxotrophs in co-culture, including two dual auxotrophs. The same general growth protocol was used as described above in the phenotype characterization, with the following modifications. Each auxotroph was initially grown in monoculture to high density in marine broth and then MBL^+Vit^. The monocultures were all normalized to OD_600_ 0.2, then combined in pairs and diluted 1:40 into MBL^-Vit^ medium. The co-cultures were grown as described above, with daily 1:40 transfers and OD_600_ measurements. As a control, the same co-cultures were grown in MBL^+Vit^ medium.

For the supernatant and lysis experiments, exponentially growing cultures of 1A01 were grown in replicate 4 mL tubes. At the collection timepoints, tubes were collected for both supernatants and lysates. For the supernatants, cells were pelleted by centrifugation (6000 g for 10 minutes at room temperature), and the supernatant carefully removed and sterilized by 0.2 εm filtration to remove any remaining cells. The supernatants were immediately frozen at −80°C. For the lysates, the cultures (containing cells suspended in supernatant) were immediately frozen at −80°C. On the day of the experiment, all lysate samples were defrosted an lysed with a microtip probe sonicator on ice (Misonix Sonicator 3000, 2 minutes total of sonication at power setting 5, in intervals of 30 seconds on/60 seconds off). The supernatants were also defrosted, and both supernatants and lysates were stored at 4°C throughout the experiment. The experiments were performed using the standard protocol described above. The supernatants and lysates were mixed in a 1:1 ratio with fresh MBL^-Vit^ media, with MBL^+Vit^ media serving as a control. The cultures were grown as described above, with daily 1:40 transfers and OD_600_ measurements for three days.

### Modeling, statistical analysis, and data visualization

The degrader and substrate parameters were based on previous work on *Vibrio.* 1A01,^80,81^ the isolate used in supernatant and lysate experiments. Vitamin affinities were measured from this work (Table 2), and approximate vitamin yields (cells per vitamin molecules) were estimated from the literature.^14,51,63^ A detailed description of the model of vitamin-limited growth on particles is available in the SI text. Numerical simulations of the model were run using the py-pde package^82^ v0.32.1 in Python v5.3.3. All statistical analyses were done using R v4.0.2^83^. Plots were created using the ggplot2 package v3.3.3^84^ and ggtree v2.4.1^85^. Genomes were visualized using clinker^86^.

## Supporting information

Table S3. Environmental vitamin measurements.

## Data availability

Metagenomic reads from seawater microcosms are accessible through NCBI BioProject (accession number PRJNA1028615).

## Code availability

Code used to generate the particle model and data analysis is available from: https://github.com/RachelGregor/marine-vitamin-auxotrophies

## Author information

R.G. and O.X.C. conceptualized this work. R.G., R.E.S., and M.G. developed the experimental methodology. R.G., R.E.S., and E.B.Q. conducted experiments. R.G. and R.E.S. analyzed experimental data. R.G., M.G., R.R., and N.M.L. performed genome analysis. G.T.V. conceptualized and executed the model. R.G. and G.T.V. created the visualizations. R.G. and O.X.C. acquired funding. O.X.C. supervised and administered the project. R.G. and G.T.V. wrote the original draft. All authors reviewed and edited the final manuscript.

## Acknowledgements

We thank all members of the Cordero group and the Simons Collaboration on Principles of Microbial Ecosystems (PriME) for enriching discussions and input, and especially to Andreas Sichert (ETH Zurich) for help with the metagenomes analysis, Annika Gomez (Columbia University) for guidance on the seawater sampling, and Stefany Moreno-Gámez for input on the modeling. We acknowledge funding from the Simons Collaboration on Principles of Microbial Ecosystems (PriME) award number 542395 (O.X.C.). R.G. is grateful for support from the Simons Postdoctoral Fellowship in Marine Microbial Ecology (653410) and the Center for Chemical Currencies of a Microbial Planet Postdoctoral Fellowship (National Science Foundation, OCE-2019589; this is the NSF Center for Chemical Currencies of a Microbial Planet (C-CoMP) publication #031).

## Extended Data

**Extended Data Fig. 1.**
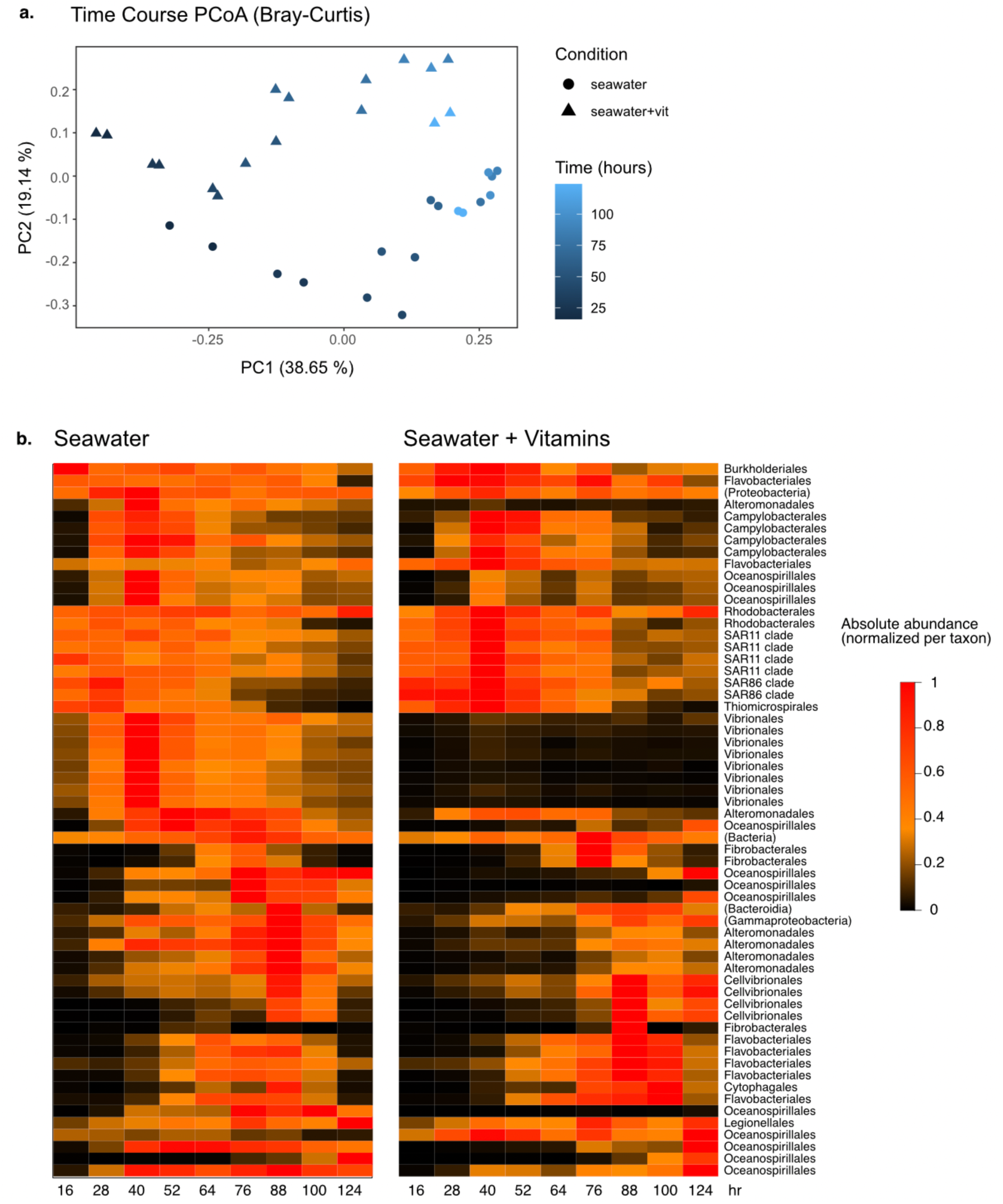
Taxonomic shifts over time in seawater microcosm experiment with and without added vitamins. A) The two conditions are distinct over time and B) differ in the abundance of key taxa. Metagenomics data was assigned taxonomy using PhyloFlash (see Methods).

**Extended Data Fig. 2.**
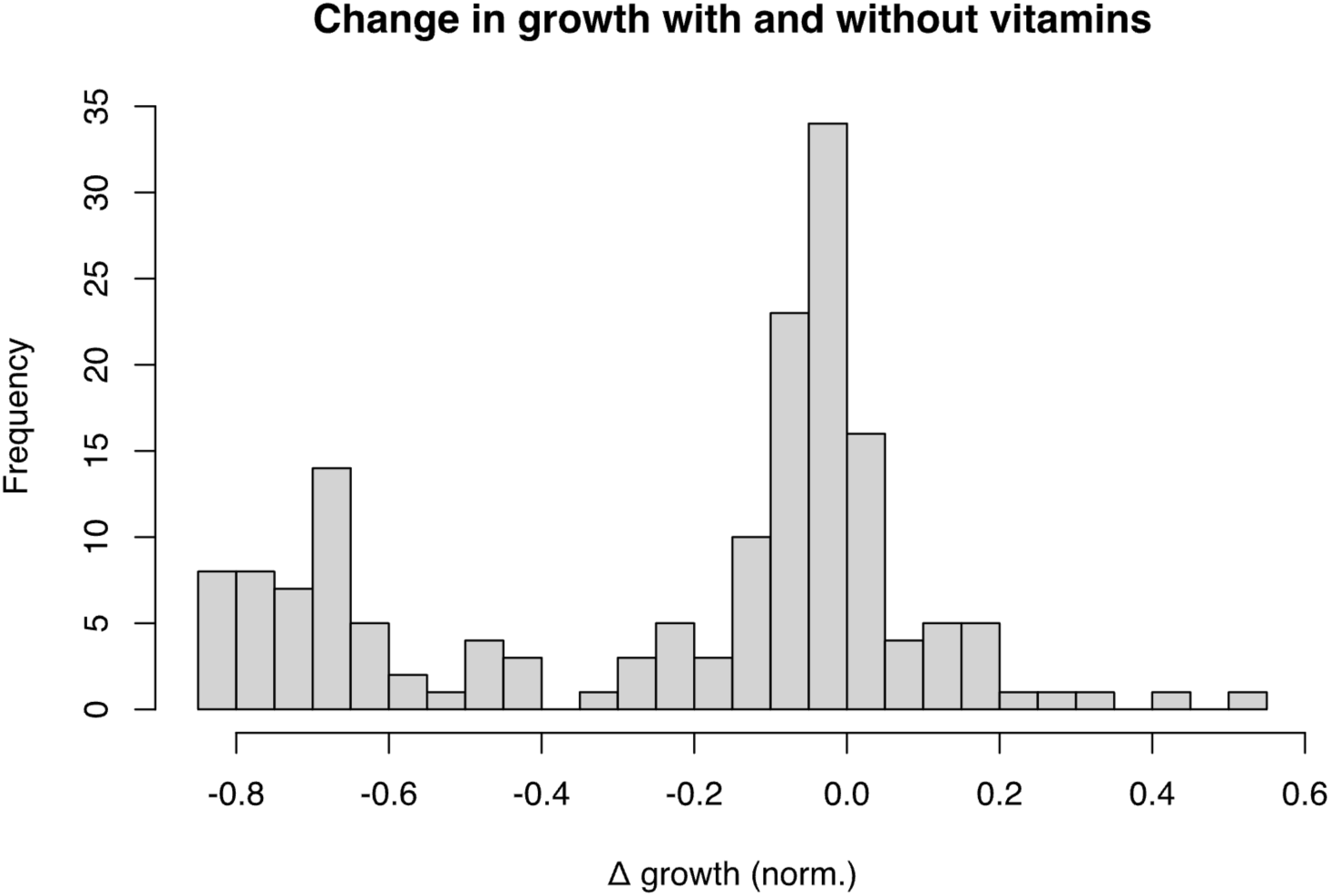
Change in growth with and without vitamins. The median OD value was calculated for both conditions and the growth deficit calculated as follows: (MBL^-vit^ -MBL^+vit^)/(MBL^-vit^ + MBL^+vit^). A growth deficit value of −0.4 was chosen as the cutoff for putative auxotrophs based on the distribution of values shown here.

**Extended Data Fig. 3.**
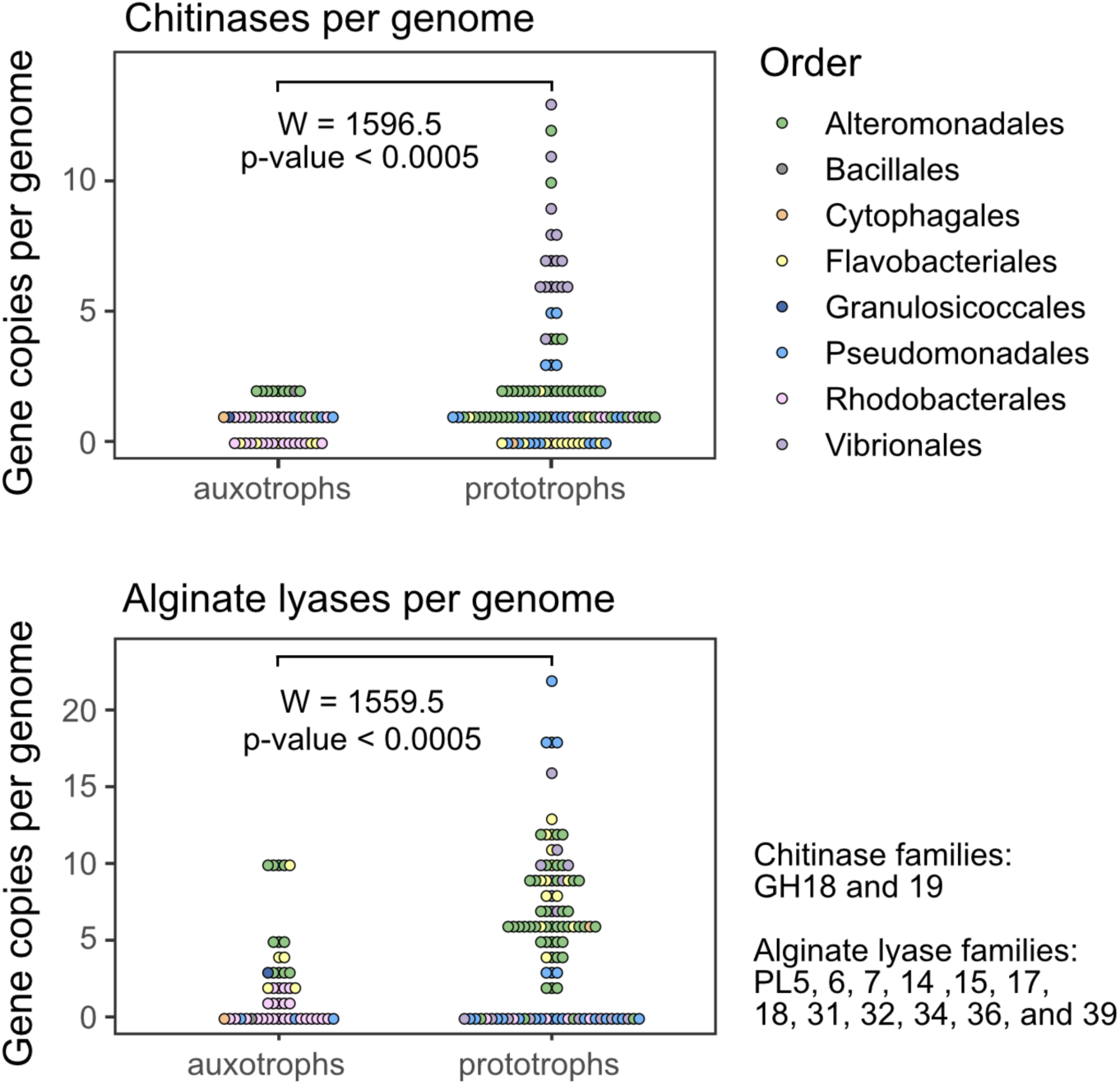
Chitinases and alginate lyases per genome. Both chitin (top) and alginate (bottom) hydrolysis genes are enriched in prototrophs compared to auxotrophs (Wilcoxon rank sum test).

**Extended Data Fig. 4.**
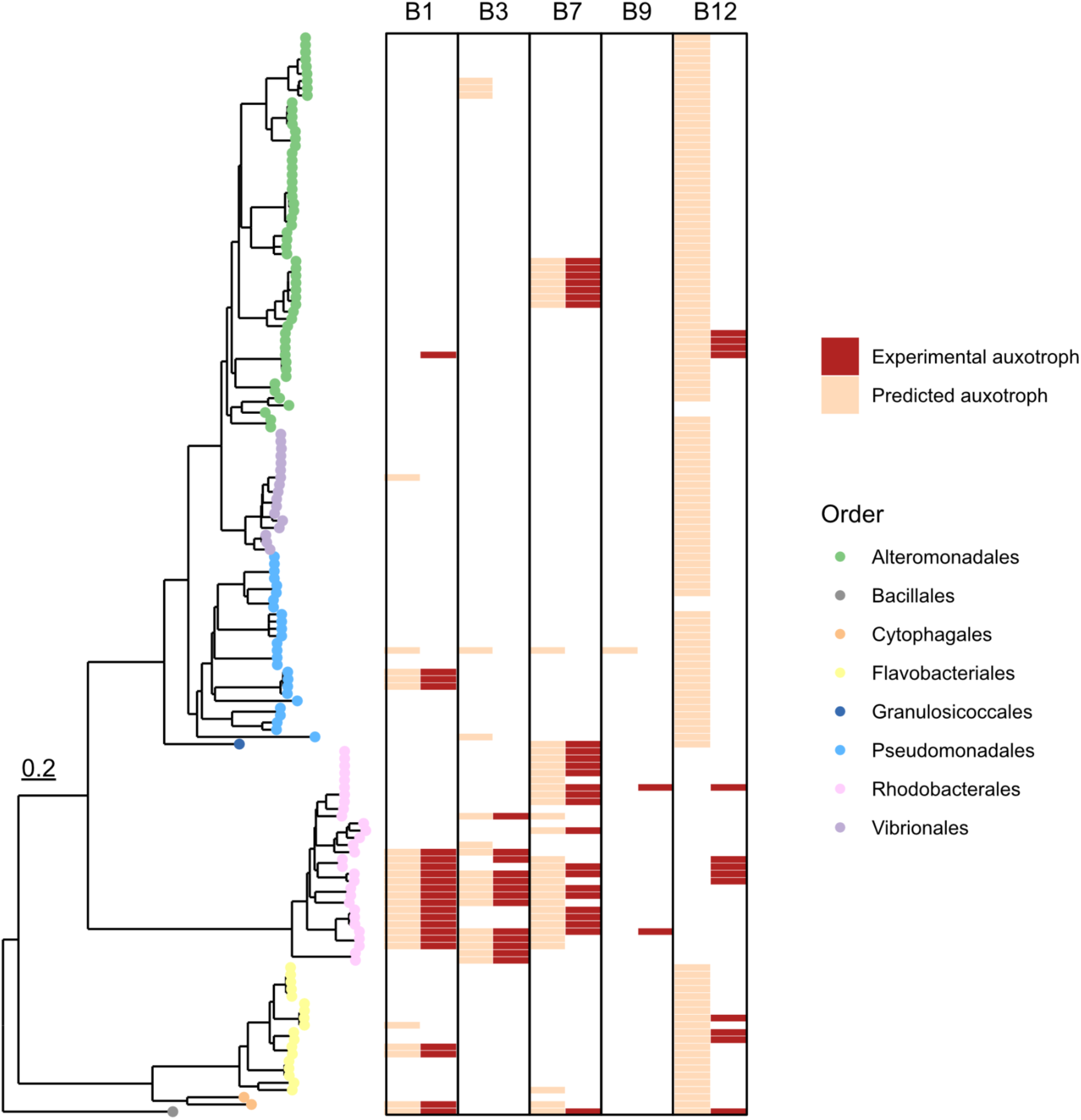
Phenotype-genotype matching based on presence/absence of key genes (see Table S1)

**Extended Data Fig. 5.**
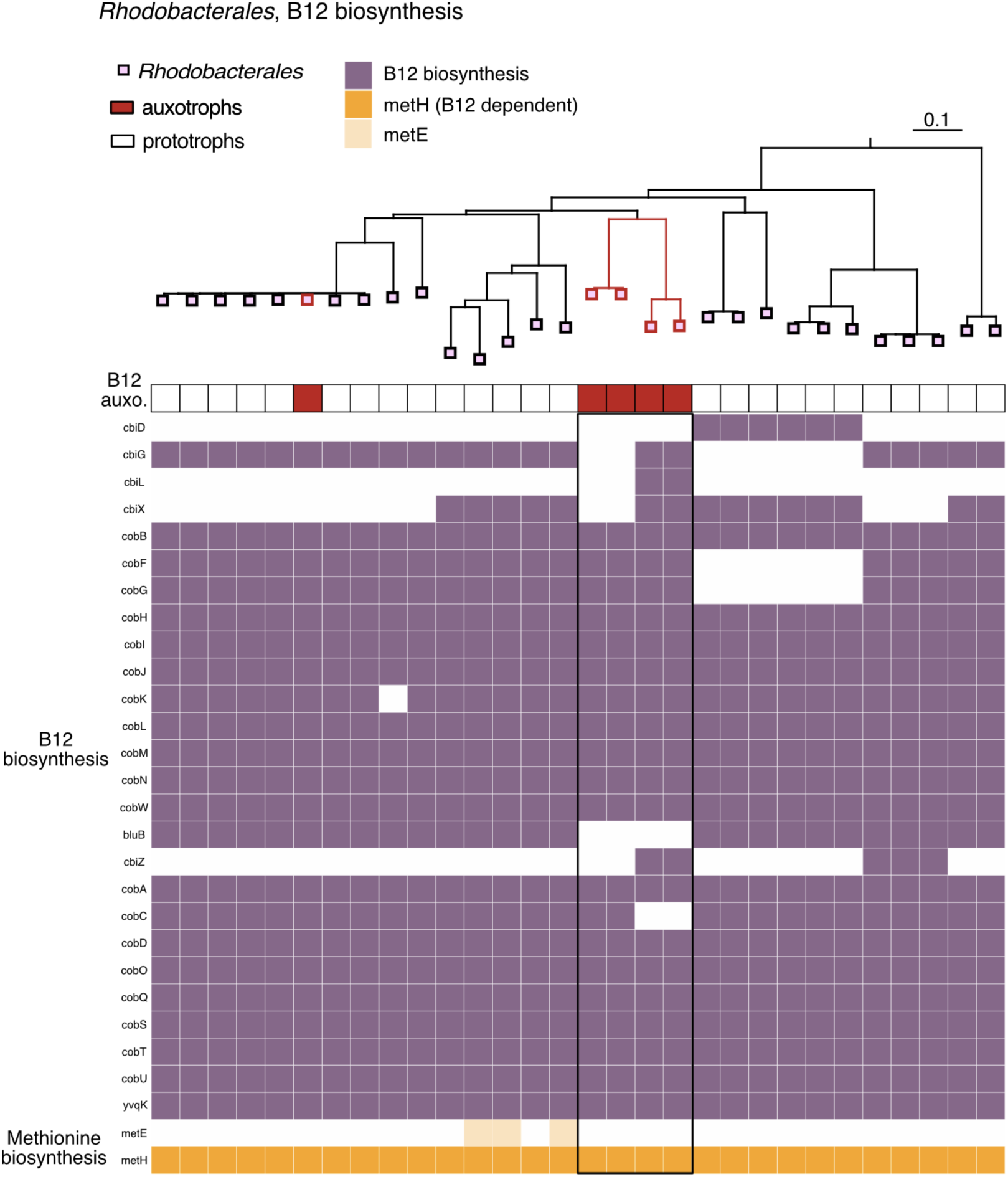
B12 biosynthesis and dependent genes in *Rhodobacterales*. All but 3 of the Rhodobacterales lack metE (peach) and are therefore dependent on B12 for methionine biosynthesis via metE (orange). The clade of four auxotrophs outlined in black are likely precursor auxotrophs for 5,6-dimethylbenzimidazole (DMB), as they are missing bluB.^87^ However, since they retain most of the pathway, they are misclassified as prototrophs based on gene predictors (Table S2).

**Extended Data Fig. 6.**
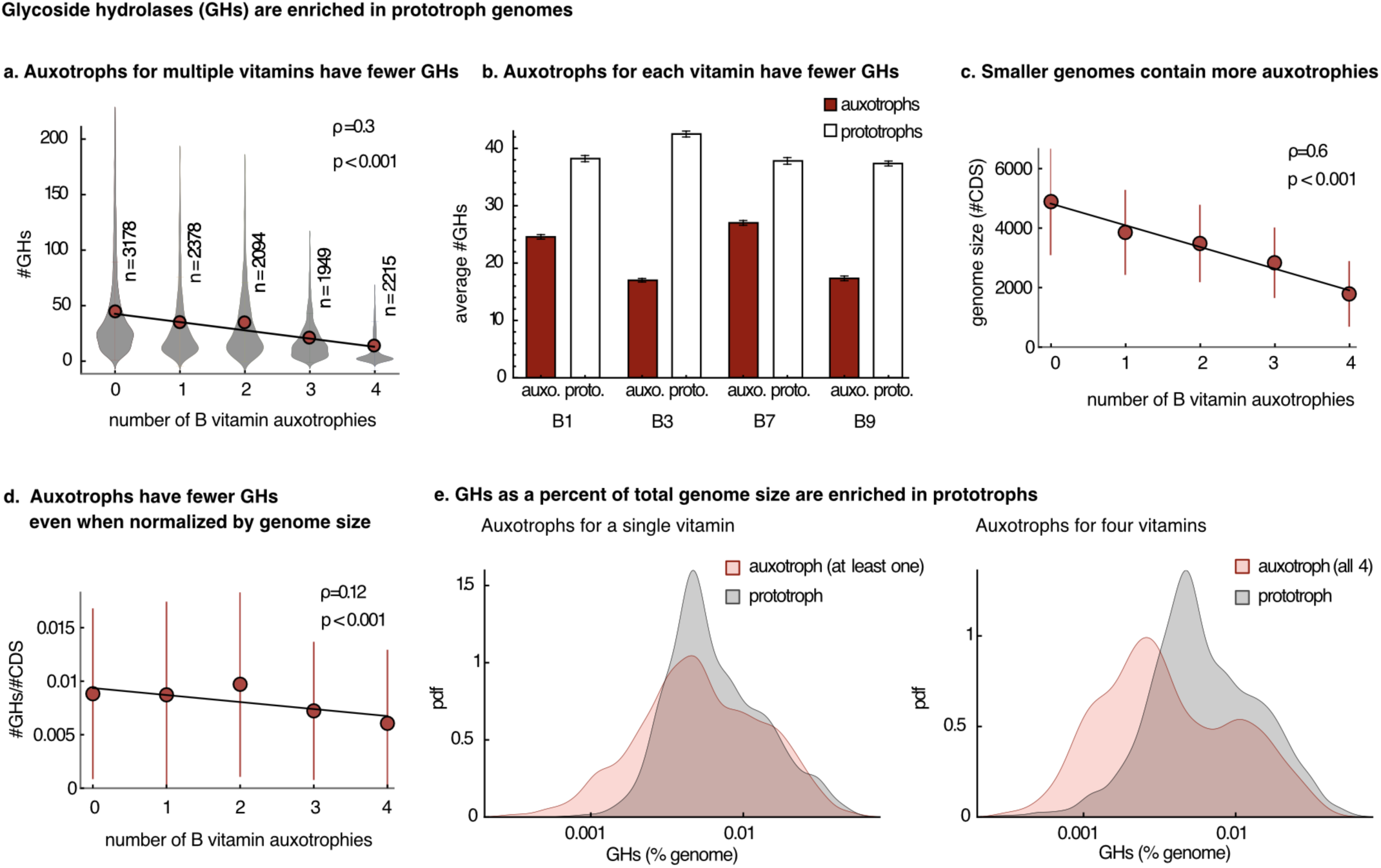
Vitamin prototrophy and polysaccharide degradation genes are positively correlated. A) We assigned auxotrophies for vitamins B1, B3, B7, and B9 based on presence-absence of key biosynthesis genes (Table S2) for 11,000 diverse reference genomes from proGenomes, and found that a class of genes for polysaccharide degradation, glycoside hydrolases (GHs), are enriched in prototrophs. When examining multiple auxotrophies in genomes, additional vitamin auxotrophies correlated to lower GH numbers on average. B) For individual vitamins, auxotrophs tended to have fewer GHs than prototrophs. C) Smaller genomes contain more auxotrophies. D) However, even when normalized by genome size (coding sequences, CDS) there is a positive correlation between the number of vitamin prototrophies and GH genes. E) GHs as a percent of total genome size are enriched in prototrophs, especially when compared to auxotrophs for all four vitamins (right-hand panel). pdf = probability density function.

**Extended Data Fig. 7.**
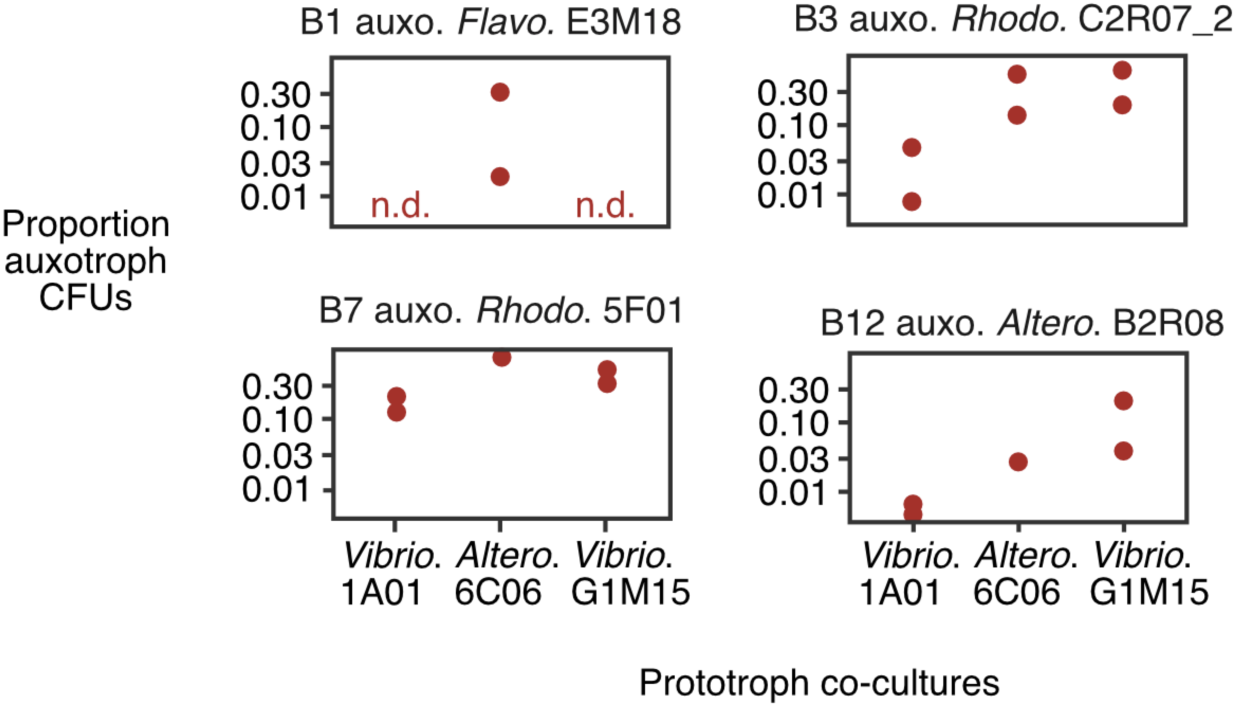
Auxotroph and prototroph co-cultures. The co-cultures were performed as in Fig. 4a. Auxotrophs and prototrophs were grown separately with vitamins, then combined 1:1 and transferred to fresh medium without vitamins. The media contained four carbon sources (glucose, glutamine, glycerol, and pyruvate) to enable the growth of isolates with different carbon preferences (see Methods). Growth was measured after 3 growth-dilution cycles. Auxotroph growth could not be directly measured by OD600, since all cultures contained prototrophs and grew to high optical densities even in the absence of vitamins. Therefore, we selected four auxotrophs that form pigmented colonies and could be differentiated from the prototrophs, and the ratio between auxotrophs and prototrophs was measured by counting colony forming units (CFUs). Cultures were measured in duplicate (two replicates were removed due to contamination for 5F01-6C06 and B2R08-6C06).

**Extended Fig. 8.**
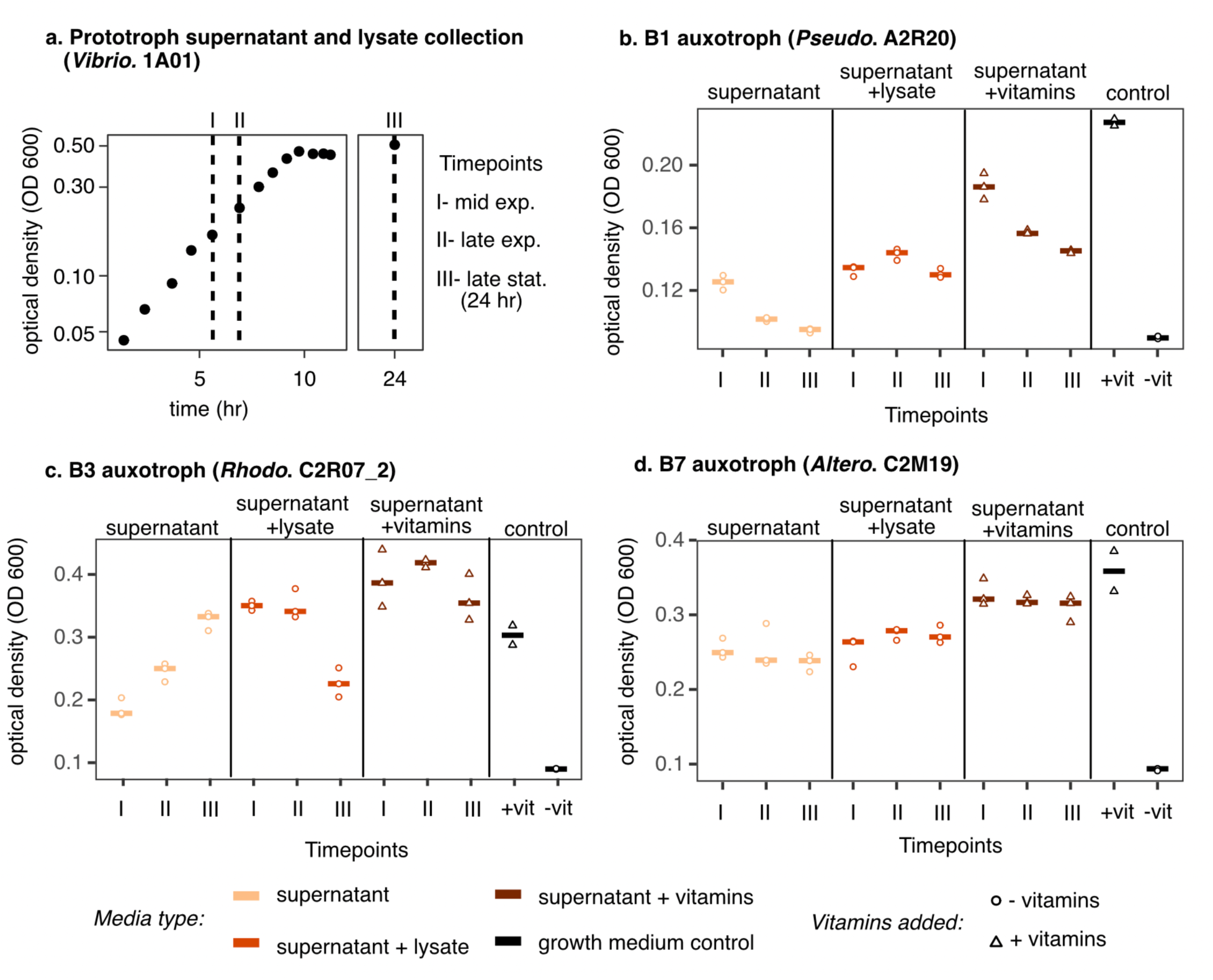
Raw data from supernatant and lysate experiment. a. Supernatants and lysates were collected from a *Vibrionales* 1A01 prototroph culture in three growth stages: mid and late exponential phase, and late stationary phase (24 hours). At each timepoint, half of the culture was used to generate sterile supernatant (peach), and the other half was harvested to generate sterile supernatant+lysate: cells were lysed in the culture using a probe sonicator, producing a mix of supernatant and lysate, which was then centrifuged and filter sterilized (red). b-d. Auxotrophs were grown on each supernatant and supernatant+lysate, as well as controls: supernatant supplemented with vitamins (brown) and media controls with and without vitamins (black). The growth of the auxotrophs was dependent on the vitamin auxotrophy as well as the growth stage of the prototroph. **Note:** In some cases, the growth on supernatants or lysates with vitamin supplementation was higher or lower than in fresh medium plus vitamins (black, control +vit), indicating that additional nutrients are being produced or depleted. To account for this, all data in Fig. 4b is normalized by the supernatant+vitamin control in each timepoint, to obtain the percentage of growth without vitamin limitation in each condition. Normalized data from timepoint II is presented in Fig. 4b.

