## Supplemental tables and text. for "Vitamin auxotrophies shape microbial community assembly in the ocean"

### Table of Contents:

I: Supplementary tables

Table S1. Auxotrophy predictions based on genomes.

Table S2. Summary of auxotrophy predictions from genomes compared to phenotypes.

Table S3. Environmental vitamin measurements from literature (see attached Excel file).

Table S4. Parameters for model of vitamin limitation on particles.

Table S5. Numerical and analytical estimates for auxotroph growth rates on particles.

II: Supplementary text: model of vitamin limitation on particles.

### I: Supplementary tables

Table S1. Auxotrophy predictions based on genomes.

| Vit. | Essential biosynthesis genes | Reference |
|---|---|---|
| **B1** | thiC and [thiG or thi4] and [thiE or thiN] | Paerl et al (2018)[13] |
| **B3** | nadC + nadD + nadE +<br>1. (nadX or nadB) + nadA or<br>2. kmo | Magnúsdóttir et al (2015)[22] and Lima et al (2009)[23]<br><br>*Note*: kmo added based on Lima et al as representative gene from kynurenine pathway for quinolinate biosynthesis via tryptophan, present in *Flavobacteriales*. |
| **B7** | bioF + bioA + bioD + bioB | Wienhausen et al (2022)[14] |
| **B9** | folK + folP | de Crécy-Lagard et al (2007)[24]<br>*Note*: While this predictor did not correctly predict auxotrophies for the experimental auxotrophs, there were only two experimental auxotrophs for B9, making it difficult to assess whether this is generalizable. Alternate predictors relying on all genes in the pathway, e.g. Magnúsdóttir et al (2015),[22] resulted in the majority of prototrophs misclassified as auxotrophs, and were therefore not used. |
| **B12** | Aerobic biosynthesis (23 total): ALA synthesis (either EC:2.3.1.37 or both EC:1.2.1.70 and EC:5.4.3.8), EC:4.2.1.24, EC:2.5.1.61, EC:4.2.1.75, EC:2.1.1.107, EC:2.1.1.130, EC: 1.14.13.83, EC:2.1.1.131, EC:2.1.1.133, EC:2.1.1.152, EC:1.3.1.54, EC:2.1.1.132, EC:5.4.99.61, EC:6.3.5.9, EC:6.6.1.2, EC:2.5.1.17, EC:6.3.5.10, EC:6.3.1.10, EC:2.7.1.156, cobinamide activation (EC:2.7.7.62), cobalamin phosphatase (EC:3.1.3.73), EC:2.7.8.26<br><br>Tetrapyrrole precursor biosynthesis (5 total): ALA synthesis (either EC:2.3.1.37 or both EC:1.2.1.70 and EC:5.4.3.8), EC:4.2.1.24, EC:2.5.1.61, EC:4.2.1.75, EC:2.1.1.107 | Shelton et al (2019)[25]<br><br>*Note*: B12 producers were classified based on this reference as very likely cobamide producers containing all 23 aerobic genes, as well as likely cobamide producers containing ≥4/5 tetrapyrrole precursor biosynthesis steps and ≥90% aerobic steps. While genes from the anaerobic pathway of B12 biosynthesis were checked as well, this pathway did not add any additional predicted producers in this dataset. |

Table S2. Summary of auxotrophy predictions from genomes compared to phenotypes.

| Vitamin | Phenotype | Genotype | Type | Frequency | % Mismatches *Per phenotype* | % Mismatches *Overall* |
|---|---|---|---|---|---|---|
| B1 | Auxo. | Auxo. | Match | 21 | Auxotrophs: 4.5% | Overall B1: 2.7% |
| | Auxo. | Proto. | Mismatch | 1 | | |
| | Proto. | Proto. | Match | 125 | Prototrophs: 2.3% | |
| | Proto. | Auxo. | Mismatch | 3 | | |
| B3 | Auxo. | Auxo. | Match | 12 | Auxotrophs: 7.7% | Overall B3: 4.7% |
| | Auxo. | Proto. | Mismatch | 1 | | |
| | Proto. | Proto. | Match | 131 | Prototrophs: 4.4% | |
| | Proto. | Auxo. | Mismatch | 6 | | |
| B7 | Auxo. | Auxo. | Match | 25 | Auxotrophs: 0% | Overall B7: 6.7% |
| | Auxo. | Proto. | Mismatch | 0 | | |
| | Proto. | Proto. | Match | 115 | Prototrophs: 8% | |
| | Proto. | Auxo. | Mismatch | 10 | | |
| B9 | Auxo. | Auxo. | Match | 0 | Auxotrophs: 100% | Overall B9: 2% |
| | Auxo. | Proto. | Mismatch | 2 | | |
| | Proto. | Proto. | Match | 147 | Prototrophs: 0.7% | |
| | Proto. | Auxo. | Mismatch | 1 | | |
| B12 | Auxo. | Auxo. | Match | 8 | Auxotrophs: 38.5% | Overall B12: 75.3% |
| | Auxo. | Proto. | Mismatch | 5 | | |
| | Proto. | Proto. | Match | 29 | Prototrophs: 78.8% | |
| | Proto. | Auxo. | Mismatch | 108 | | |

Table S3. Environmental vitamin measurements from literature (see Excel file).

Table S4. Parameters for model of vitamin limitation on particles.

| Parameter name | Value | Reference or formula |
|---|---|---|
| Cross-feeder maximum growth rate ($r_{cf}$) | $r_{cf} = 0.3h^{-1}1$ | Gralka et al (2023)[21] |
| Yield on acetate ($Y_{act}$) | $31\ OD/M_{act}$ | Value measured for marine isolate 3B05 growing on acetate[81] |
| CFU/ml to OD conversion | $9 \cdot 10^8 \frac{CFU/ml}{OD}$ | Value measured for marine isolate 1A01 growing on glucose[81] |
| Yield on vitamins (molecules per cell) | 20,000 (B1); 100,000 (B3); 100 (B7); 5,000 (B12) | Approximated based on averages of published measurements[14,51,63] |
| Affinity on acetate ($K_{s/m}$) | 1 mM | General order of magnitude taken from E. coli[88–90] |
| Affinity on vitamins ($K_{s/v}$) | Variable | Measured in this work (Table 2) |
| Diffusion constant of substrate ($D_M$) | $12 * 10^{-6} cm^2/s$ | The diffusion constant for acetate was used in all simulations |
| Diffusion constant of vitamins ($D_V$) | $3.67 * 10^{-6} cm^2/s$ | The diffusion constant for B12 was used in all simulations |
| Degrader lysis rate ($\mu$) | $\mu = 0 - 0.26h^{-1}$ | Free parameter |
| Degrader growth rate ($r$) | $r = 0.26h^{-1}$ | Guessous et al (2023)[80] |
| Equilibrium density of degraders with $\mu = 0$ ($\sigma_0$) | $7\ cells/\mu m^2$ | $\sigma_0 = \rho_b \eta \frac{V}{N_c 4\pi R_0^2}$ from Guessous et al (2023)[80] |
| Density of a monolayer of cells ($\rho$) | $0.5\ cells/\mu m^3$ | Estimated from approximate cell volume of $2\ \mu m^3$ |
| Equilibrium density of degraders ($\sigma_{eq}$) | $\sigma_{eq} = 0 - 7\ cells/\mu m^2$ | $\sigma_{eq} = \left(1 - \frac{\mu}{r-d}\right)\sigma_0$ |
| Specific activity of chitinases ($k_e$) | $24\ nmol_{Gln}/\mu g_e/h$ | Guessous et al (2023)[80] |
| Enzyme per biomass on particle surface ($Y_e{}^{-1}$) | $25\ \mu g_e/OD/ml$ | Guessous et al (2023)[80] |
| Acetate produced per GlcNAc consumed ($f_{act/gln}$) | $1.48\ mol_{Act}/mol_{Gln}$ | Flux balance analysis (FBA) for 1A01[91] |
| Flux of acetate from particle surface per cell density ($F_{act}/\sigma_{eq}$) | $1.65 \cdot 10^5 \frac{molecules/s}{CFU}$ | $\frac{k_e f_{act/gln}}{Y_e}$ |
| Flux of vitamins from particle surface ($F_{vit}$) | Variable | $F_{vit} = \frac{\sigma_{eq}}{Y_{vit}}\mu$ |

Table S5. Numerical and analytical estimates for auxotroph growth rates on particles.

| **Strain (auxo.)** | **Analytical approximation of $I$** | **Analytical growth percentage ($r_{auxo}/r_{max}$)** | **Numerical estimation of $I$** | **Numerical growth percentage ($r_{auxo}/r_{max}$)** |
|---|---|---|---|---|
| A2R20 (B1) | 0.180 | 15.25% | 0.179 | 15.18% |
| E3M18 (B1) | 10.1 | 90.99% | 9.94 | 90.85% |
| B2R09 (B3) | 0.00187 | 0.187% | 0.00187 | 0.187% |
| C2R07_2 (B3) | 0.233 | 18.90% | 0.232 | 18.83% |
| 5F01 (B7) | 0.0663 | 6.22% | 0.0662 | 6.21% |
| C2M19 (B7) | 0.0278 | 2.70% | 0.0278 | 2.70% |
| B2R08 (B12) | 15.4 | 93.90% | 15.1 | 93.79% |

**II: Supplementary text: model of vitamin limitation on particles.**

To put the measured vitamin auxotrophies in an ecological context, we develop a simplified model of chitin particle degradation in the ocean. The focus of the model is to understand to what extent lysis of prototrophic degrader strains can support the growth of auxotrophic cross-feeders through vitamin sharing. In the following sections, we will first describe the biological parameters in the model controlling degrader and cross-feeder growth. Then, we will move on to the physical parameters describing the geometry of the chitin particle and the spatial dynamics of vitamins and cross-fed metabolites. Lastly, we will provide details about the numerical simulations performed and develop approximate analytical solutions to model equations. It is important to note that in this model we assume the timescale of diffusion processes is much faster than the timescale of microbial growth. Therefore, we use the steady-state solution for all diffusion processes described to calculate biological parameters like growth rates.

**Biological parameters:**

We model the growth dynamics of degrader growth on a particle as a logistic growth with an extra death term representing lysis:

$$\frac{d\sigma}{dt} = r\sigma\left(1 - \frac{\sigma}{k}\right) - \mu\sigma$$

In this equation, $\sigma$ represents the density of degraders on the particle and $k$ the carrying capacity of the particle, both with units of $cells/um^2$. The degrader growth rate is represented by $r$ and the lysis rate is represented by $\mu$, both with units of $h^{-1}$. For the purposes of our model, we assume the degrader population to be at steady-state and therefore not changing with time. Calculating this steady-state density, we arrive at:

$$\sigma^* = \left(1 - \frac{\mu}{r}\right)k$$

We parametrize this equation with previously measured values for carrying capacity and growth rate for the 1A01 strain, a model chitin degrader.[80] For this strain, the growth rate on chitin particles has been measured to be $0.26\ h^{-1}$ and we estimate the carrying capacity of chitin particles to be about $7\ cells/um^2$ from optical density data collected during the stationary growth phase of 1A01 liquid cultures.[80] Adopting these numbers, the steady-state density of degraders on the particle becomes only a function of the lysis rate and it goes to zero as the lysis rate approaches the degrader growth rate.

Having established the steady-state dynamics of degrader growth and death on the particle, we now consider the ability of cross-feeders to grow in this environment. We assume that the growth of auxotrophic cross-feeders depends on the release of both substrates (for example metabolic byproducts like acetate) and vitamins by degraders. We model the cross-feeder growth rate as a multiplicative Monod function of the concentration of available substrates and vitamins:

$$r_{auxo} = r_{cf}\left(\frac{[M]}{[M]+K_{s,m}}\right)\left(\frac{[V]}{[V]+K_{s,v}}\right)$$

Where $M$ represents the concentration of substrates, $V$ represents the concentration of vitamins, and $K_{s,m}, K_{s,v}$ represent their respective Monod constants. In addition, $r_{cf}$ represents the maximum growth rate of the cross-feeder strain when neither substrates nor vitamins are limiting. We further define the maximum growth rate of the cross-feeder strain for a given substrate concentration:

$$r_{max} = r_{cf}\left(\frac{[M]}{[M]+K_{s,m}}\right)$$

We parametrize these equations with literature values and quantities measured in this study. We take the value for the maximum growth rate of cross-feeders to be $0.3\ h^{-1}$ as estimated in a large screen previously conducted in our lab with the same isolate collection used in this study.[21] We focus on acetate, which is secreted by 1A01 during growth and has been previously shown to support cross-feeding.[81] We use a value of $K_{s,m} = 1\ mM$ based on previous studies of the growth of *E. coli* on acetate.[88–90] For vitamins, we use the Monod constants estimated in this study.

**Physical parameters:**

For this simplified model, we assume a radially symmetric, spherical chitin particle with radius $R_0 = 150\ \mu m$. Both degraders and cross-feeders colonize the particle surface. As the degraders break down the particle and grow, they produce substrates and vitamins that are consumed by the cross-feeders. However, these molecules simultaneously freely diffuse from the surface of the particle into the ocean while they are being consumed, and we model these processes with the following equations:

$$\frac{\partial M}{\partial t} = D_M \nabla^2 M - r_{auxo}\frac{\rho}{Y_M}$$

$$\frac{\partial V}{\partial t} = D_V \nabla^2 V - r_{auxo}\frac{\rho}{Y_V}$$

Where $M$ and $V$ represents the concentrations of metabolic byproducts and vitamins, respectively, $D_M$ and $D_V$ their diffusion constants, and $Y_M$ and $Y_V$ the yield of cells on those resources. Additionally, $r_{auxo}$ is the auxotroph growth rate defined previously and $\rho$ is the volume density of cross-feeder cells. Focusing on the initial moments of cross-feeder colonization of the particle, we assume $\rho$ to be a $1\mu m$ thick monolayer of cells and set the cross-feeder density to 0 beyond that point.

To fully define these equations, we are missing boundary conditions at the surface of the particle and at infinity. For the purposes of our model, we assume that the ocean contains no substrates or vitamins, making their concentration approach zero as we move away from the particle. At the surface of the particle, however, there is a flux of both substrates and vitamins determined by the growth and lysis of degrader cells.

To calculate the flux of substrates, we first calculate the rate of chitin degradation and the release of its monomer, N-acetyl glucosamine (GlcNAc) as the product of the cell density on the particle surface ($\sigma *$), the conversion factor between chitinase mass and cellular biomass ($Y_e{}^{-1}$), and the chitinase specific activity ($k_e$). All these values were experimentally determined for 1A01 by Guessous and co-authors.[80] We further make the simplifying assumption suggested by Guessous and co-authors that 1A01 is capable of uptaking all the GlcNAc produced during particle degradation, making the GlcNAc production rate equal to the GlcNAc consumption rate at the particle surface.[80] Finally, we multiply this consumption rate by how many moles of substrates are produced by 1A01 for each mole of GlcNAc consumed, arriving at the rate of substrate production on the particle surface. As an approximation for this metabolic conversion factor, we consider the ratio of moles of acetate produced to moles of GlcNAc consumed by the degrader ($f_{Act/Gln}$), which was estimated in a recently developed flux balance analysis model for 1A01.[91] Overall, the final formula for the flux of byproducts on the particle surface becomes:

$$F_M = \frac{\sigma^*}{Y_e} k_e f_{Act/Gln}$$

To calculate the flux of vitamins, we simply multiply the cell density on the particle surface ($\sigma *$), the amount of vitamins per cell ($Y_V{}^{-1}$), and the lysis rate ($\mu$). The final equation for this flux then becomes:

$$F_V = \frac{\sigma^*}{Y_V} \mu$$

The values used for each of these numerical parameters can be found in Table S4.

**Numerical simulations:**

We simulate these equations with the parameters from Table S4 using the py-pde package v0.32.1 in Python v5.3.3. With this package, we create an extension of their standard PDEBase code to include the reaction-diffusion terms shown before. For the simulation space, we use a spherically symmetric grid, no flux boundary conditions on the particle surface, and the following Robin boundary conditions at the opposite end of the simulation grid to account for finite dimensions:

$$\partial f(R_f) + \frac{1}{R_f} f(R_f) = \frac{f(\infty)}{R_f}$$

Where $f(R)$ represents the concentration of substrates or vitamins at a certain distance $R$ from the particle, $R_f$ is the distance from the center of the particle to the outside boundary of the simulation and $f(\infty)$ is the concentration of substrates or vitamins at infinity. This identity can be derived because there are no sources or sinks outside of the simulation grid, so the diffusion-reaction equations can be simplified and analytically solved in that region to be of the form:

$$f(R) = A - \frac{B}{R} \text{ for } R > R_f$$

Where A and B are numerical constants and A equals $f(\infty)$. Applying this analytical formula at the boundary of the simulation, we can calculate the value of $f$ and its derivative at $r = R_f$:

$$f(R_f) = A - \frac{B}{R_f}, \frac{\partial f}{\partial R}|_{R_f} = \frac{B}{R_f^2}$$

Plugging these values back into the expression for our Robin boundary condition verifies its validity. In our simulations, we set $R_f = 160\ \mu m$, making our simulation grid $10\ \mu m$ wide, and $f(\infty) = 0$ since we assume no vitamins or substrates are present in the ocean. We used a spatial discretization length of $0.33\ \mu m$ yielding 30 discrete simulation points.

To numerically estimate the steady state profiles of substrates and vitamins, we initialized their spatial distributions with analytical solutions to the standard diffusion equations obtained by disregarding consumption terms. We then simulated the temporal dynamics of the full reaction-diffusion equation for a period of time 100 times longer than it would take for diffusion alone to homogenize the length of our simulation space:

$$t_{sim} = 100 \cdot t_{diff} = 100 \cdot \frac{1}{2}\frac{L^2}{D}$$

Where $L = R_f - R_0 = 10\ \mu m$ is the length of the simulation grid and $D = min(D_M, D_V)$ is the minimum between the vitamin and substrate diffusion constants. We verified that longer simulation times and initialization of the profiles with the concentrations set to zero everywhere did not change the results of these numerical estimates.

**Analytical approximations:**

In addition to the numerical simulations described previously, we can derive approximate analytical expressions to gain more insight into how each factor in the model contributes to the final results. The main approximation we make is that the non-linear consumption term in the reaction-diffusion equation for vitamins is negligible. This is a reasonable approximation in our case because consumption of vitamins only occurs very close to the surface of the particle and at a small rate compared to the rate of loss to diffusion. We also numerically confirmed the validity of this approximation for each of the strains for which affinities were measured. This way, we assume the dynamics of vitamins is dominated by diffusion and solve for the vitamin concentration profile ($V$) as a function of the radial distance from the particle ($R$).

$$\boldsymbol{V(R) = \frac{{R_0}^2 F_V}{D_V R} = \frac{{R_0}^2 \sigma^* \mu}{Y_V D_V R}}$$

Where, as defined previously, $R_0$ is the radius of the chitin particle, $F_v$ is the flux of vitamins at the particle surface, and $D_v$ is the diffusion constant for vitamins. From this analytical profile, we can calculate the vitamin concentration at the particle surface to be $V(R_0) = \frac{R_0 \sigma^* \mu}{Y_V D_V} \propto {Y_v}^{-1}$ and get an expression for the growth rate of auxotrophic cross-feeders:

$$r_{auxo} = r_{max}\left(\frac{V(R_0)}{V(R_0) + K_{s,v}}\right) = r_{max}\left(\frac{V(R_0)/K_{s,v}}{V(R_0)/K_{s,v} + 1}\right)$$

Looking at this expression, it becomes clear that we can define an index $I$ that determines the qualitative outcome of growth as follows:

$$I = V(R_0)/K_{s,v} \;\propto\; {Y_v}^{-1}{K_{s,v}}^{-1}$$

If $I \gg 1$, the growth rate is high, and if $I \ll 1$, it is low. The calculations of this quantity for each of the strains for which affinities were measured are summarized in Table S5 (assuming an optimal lysis rate of $0.13\ h^{-1}$).
